# Abundance, identity, and potential diazotrophic activity of *nifH*-containing organisms at marine cold seeps

**DOI:** 10.1101/2024.07.31.606060

**Authors:** Amanda C. Semler, Emily R. Paris, Anne E. Dekas

## Abstract

Diazotrophic microorganisms alleviate nitrogen limitation at marine cold seeps using nitrogenase, encoded in part by the gene *nifH*. Here, we investigated *nifH*-containing organisms (NCOs) inside and outside six biogeochemically heterogeneous seeps using amplicon sequencing and quantitative real-time PCR (qPCR) of *nifH* genes and transcripts. We detected *nifH* genes affiliated with anaerobic methane-oxidizing ANME-2 archaea and sulfate-reducing Desulfobacteraceae, consistent with previous studies, but also phylogenetically and likely metabolically diverse organisms, including *Desulfoglaeba*, *Candidatus* Methanoliparia, and Desulfuromonadales. In total, we recovered 10,734 bona fide *nifH* sequence variants affiliated with 18 bacterial and archaeal phyla (17 within seeps), a subset of which were transcribed at nearly all seeps investigated. We corrected our qPCR data based on our amplicon results, which found that 71% of recovered sequences were not bona fide *nifH,* and we recommend a similar correction in future qPCR studies that use broad *nifH* primers. *NifH* abundance was up to three orders of magnitude higher within seeps, was highly correlated with *mcrA* gene abundance, and, when corrected, was negatively correlated with porewater ammonium <25 uM, consistent with the inhibition of diazotrophy by ammonium. Our findings significantly expand the known diversity of NCOs at seeps and emphasize seeps as hotspots for deep-sea diazotrophy.

## Introduction

Cold seeps are seafloor environments in which methane and other hydrocarbons emerge from the sediment at high concentrations, sustaining rich communities of microorganisms with a diverse array of metabolisms (Knittel & Boetius, 2009; Ruff et al., 2015). Microbial communities at seeps consume the majority of methane before it reaches the water column (Reeburgh, 2007), reducing natural emissions of this greenhouse gas and highlighting the importance of understanding the biogeochemical controls on microbial activity at seeps. Seeping hydrocarbons introduce high amounts of carbon, but not nitrogen, to sediments, which can lead to nitrogen depletion within cold seep ecosystems and potentially limit overall productivity (Dekas et al., 2009; Joye, 2020; Joye et al., 2004). Biological nitrogen fixation (diazotrophy) is one way in which microorganisms can produce their own bioavailable nitrogen. In this process, nitrogen gas is converted to ammonia, with the critical step – mediated by the enzyme nitrogenase – breaking the N_2_ triple bond.

The most widely sequenced marker gene for nitrogenase, *nifH*, has been found at cold seeps around the world, primarily using clone libraries and qPCR (Dang et al., 2009; Dekas et al., 2009, 2014, 2016, 2018; Miyazaki et al., 2009), but also in metagenomic surveys (Dong et al., 2022; Zhao et al., 2020). Furthermore, nitrogen fixation has been detected at multiple deep-sea cold seeps via uptake of ^15^N_2_ (Dekas et al., 2009, 2014, 2016, 2018; Metcalfe et al., 2021). Diazotrophy has only been experimentally validated in the anaerobic methanotrophic ANME-2 archaea and their sulfate-reducing bacterial symbionts (Dekas et al., 2009, 2014, 2016; Metcalfe et al., 2021), but the *nifH* sequences recovered at seeps span a wide range of *nifH* phylogeny (groups I, II, III, and beyond), suggesting there is a taxonomically diverse community of diazotrophs. Phylogenetically identifying *nifH*-containing organisms (NCOs) based on *nifH* sequences is challenging and prone to inaccuracies, due to horizontal gene transfer events and the resulting incongruence of ribosomal and *nifH* gene trees (Kapili & Dekas, 2021). As a result, the taxonomic identity of many NCOs remains largely unknown, and their overall ecological role at seeps unconstrained.

One recent study, which investigated metagenomes from 11 cold seeps, identified NCOs by detecting *nifH* genes within metagenome-assembled genomes (MAGs). They identified *nifH* within ten different phyla: Halobacteriota (including ANME), Desulfobacterota, Chloroflexota, Verrucomicrobia, Firmicutes, and five candidate phyla not previously known to host *nifH* genes: Altarchaeota, Omnitrophota, FCPU426, Caldatribacteriota and UBA6262, along with genes for diverse inorganic metabolisms and heterotrophy (Dong et al., 2022). Remarkably, this diversity was captured in just 35 *nifH*-containing MAGs, which represented 4% of the total microbial communities. Gene abundance ratios comparing *nifH* to single-copy ribosomal protein genes on the assembled contigs from these sites indicated NCOs comprised 24% of the total microorganisms surveyed, suggesting the true diversity of diazotrophs could be much greater than that represented in the MAGs (Dong et al., 2022). Indeed, the MAGs only included about half of the unique *nifH* sequences (n = 66) detected in the assembled contigs from those sites.

In parallel with the taxonomic diversity of *nifH*, previous studies have demonstrated high concentrations of *nifH* genes at seeps. Two previous qPCR surveys of the South China Sea (Dang et al., 2009; n = 6 seeps) and of Kumano Knoll (Miyazaki et al., 2009; n = 1 seep) both used *nifH* primers [mehtaFw-28/mehtaRv-416; (Mehta et al., 2003)] to determine *nifH* abundances of roughly 10^6^ – 10^8^ copies per gram wet sediment in seep sediments. However, *nifH* primer sets, including the widely used mehtaFw-28/mehtaRv-416 set, tend to be massively degenerate in order to detect the full phylogenetic diversity of *nifH* sequences in the environment (Gaby & Buckley, 2012). As a result, they also commonly detect homologs of *nifH* not involved in nitrogen fixation, referred to as “*nifH*-like” sequences. These various *nifH* groups (IV-VI) include group IV-D – containing methanogenesis cofactor F430 biosynthesis genes *cfbC* and *cfbD* – and group V – containing bacteriochlorophyll and chlorophyll biosynthesis genes *bchX*, *bchL*, and *chlL* (Dong et al., 2022; Gaby & Buckley, 2012; Ghebreamlak & Mansoorabadi, 2020; Staples et al., 2007). Recent studies have shown that these *nifH*-like sequences can represent more than half of the sequencing reads discovered in marine sediments using the mehtaFw-28/mehtaRv-416 primers, and that the proportion can vary by habitat and sample type (Kapili et al., 2020). Although this is not a problem for sequencing studies aiming to explore genes related to nitrogen fixation, which can filter out *nifH*-like reads, it is more problematic for qPCR applications. It remains undetermined how much this overestimate affects cold seep environments, and therefore whether the previous *nifH* quantifications at seeps are robust.

In the present study, we sequence and quantify *nifH* genes and transcripts within sediments collected inside and outside six cold seep sites. The seeps investigated are from two geologically distinct regions, the U.S. Atlantic continental margin (USAM) and Monterey Bay, California (MB), and they encompass significant physicochemical and biological diversity. While the USAM seep system contains relatively classic cold seep chemistries and microbial communities (Semler et al., 2022), the MB seeps contain non-methane hydrocarbons in addition to methane (Lorenson et al., 2002; Orange et al., 1999; Stakes et al., 1999) and atypical seep microbial communities (Semler & Dekas, 2024, preprint). In particular, the MB seeps virtually lack ANME-2 archaea, potentially creating a need for other diazotrophs. We combine a highly spatially resolved sampling scheme with deep amplicon sequencing of *nifH* genes and transcripts to recover orders of magnitude more *nifH* sequences per sample and site than previous studies. Then we use a recently developed pipeline to identify bona fide *nifH* sequences and infer *nifH*-host taxonomy in a high throughput fashion, improving upon previous BLAST-based approaches by using both sequence identity and phylogenetic approaches (Kapili et al., 2020). Finally, we perform qPCR, corrected using a sample-specific reduction factor to exclude *nifH*-like sequences, and thus quantify NCOs throughout these systems. We compare their abundance with geochemical (CH_4_ ^+^, S^2-^, NH_4_ ^+^, NO_2_ ^-^, and NO_3_ ^-^) and biological (*mcrA* gene abundance) parameters. Together, our study presents the abundance, distribution, taxonomic affiliation, and potential diazotrophic activity of NCOs at marine cold seeps.

## Methods

### Site description, sample collection, and processing

We collected sediment cores from inside and outside six cold seep sites – Shallop Canyon East (USAM-SE), Shallop Canyon West (USAM-SW), New England (USAM-NE), and Veatch Canyon (USAM-VC) on the U.S. Atlantic Margin; and Clam Field (MB-CF) and Extrovert Cliff (MB-EC) in Monterey Bay – which ranged from 335 to 1,545 mbsl (Fig. S1). Core retrieval and on-board processing are detailed in Semler et al. (2022) (USAM sites) and in Semler & Dekas (2024, preprint; MB sites). Briefly, sediment pushcores were collected from inside each cold seep (hereafter called “Seep” cores; n = 10 cores) and in nearby background sediments (5-500 meters from visual indicators of seepage; hereafter called “Background” cores; n = 12 cores). Cores were up to 20 cm in length and were sectioned into 3 cm (USAM) or 2.5 or 5 cm (MB) horizons. Sediment core information is listed in Table S1. Several 1 mL subsamples of each depth horizon were immediately flash-frozen in liquid nitrogen and preserved at - 80 °C for later DNA and RNA extraction. Porewater was also collected from sediments immediately after sectioning using either Rhizon samplers inserted through pre-drilled holes (USAM) or a porewater pressing bench (KC Denmark Research Equipment, Silkeborg, Denmark) under a stream of argon gas (MB). Porewater was filtered with 0.2 µm Durapore® PVDF membrane filters (EMD Millipore, Burlington, MA, USA).

### Geochemical measurements

Concentrations of ammonium, nitrite, and nitrate in MB seep and background sediments were measured and are reported here for the first time. Ammonium concentrations were determined colorimetrically from porewater using the indophenol blue method (Bower & Holm-Hansen, 1980), with a detection limit of 0.5 µM. Nitrite and nitrate (NOx – nitrite) concentrations were determined colorimetrically using the vanadium(III) chloride method (Schnetger & Lehners, 2014), with detection limits of 0.1 µM and 1 µM, respectively. Concentrations of other geochemical variables at MB sites, including methane and sulfide, were reported in Semler & Dekas (2024, preprint). Concentrations of all these geochemical variables were previously reported for USAM sites in Semler et al. (2022).

### Nucleic acid extraction, amplification, and sequencing of nifH genes and transcripts

DNA and RNA were extracted from flash-frozen sediments using the RNeasy Powersoil Total RNA Isolation Kit (RNA) and the RNA Powersoil DNA Elution Accessory Kit (DNA; MoBio Laboratories, Carlsbad, CA, USA) from sediments flash-frozen on board and stored at −80 °C. RNA extracts were cleaned with the Ambion TURBO DNA-free Kit (ThermoFisher Scientific, Waltham, MA, USA), and reverse transcription of RNA to cDNA was completed using Superscript III First Strand Synthesis Supermix (Thermofisher Scientific, Waltham, MA, USA). Lack of DNA contamination in the RNA extracts was confirmed by processing RNA extracts (without reverse transcription) in parallel and seeing no visible amplification on a gel after the second PCR.

DNA and cDNA were concentration normalized and amplified using a two-step PCR plan for Illumina amplicon sequencing. In the first step, *nifH* primers mehtaFw-28/mehtaRv-416 (Mehta et al., 2003) were used to target *nifH* genes and transcripts. The gene-targeting region of this primer set is listed in Table S2. Both sets of primers included an extension complimentary to the primers used in the second PCR. 25 µL PCR reactions were performed containing 0.5 µL of forward and 0.5 µL of reverse primers (10 µM concentration), 12.5 µL Takara ExTaq (2x, Quanta-Bio, Beverly, MA, USA), 0.5 µL 2.5 mg/mL bovine serum albumin, 10 µL DNase-free water, and 1 µL DNA or cDNA template. The thermal cycling conditions were as follows: initial denaturing at 95 °C for 120 s; 40 cycles of 95 °C for 30 s, 55 °C for 30 s, and 72 °C for 45 s; a final elongation step at 72 °C for 300 s; and refrigeration at 4 °C until removal and storage.

In the second step, Illumina adaptors, barcodes, and indices were added to the amplicons. The same PCR reaction mix was used with custom primers targeting the primer extension in the first PCR. The thermal cycling conditions were as follows: initial denaturing at 95 °C for 180 s; 8 cycles of 95 °C for 30 s, 55 °C for 30 s, and 72 °C for 30 s; a final elongation step at 72 °C for 300 s; and refrigeration at 4°C until removal and storage. Amplicons were cleaned with 0.7x AMPure XP magnetic beads (Beckman-Coulter, Brea, CA, USA), pooled, and quantified before being sent to the UC Davis DNA Technologies Core Facility (Davis, CA, USA) for Illumina MiSeq 2×250 bp sequencing. Seventeen samples were randomly chosen for duplicate amplification, and the average weighted UniFrac distance between duplicate samples was 0.153. Positive controls of *nifH* mock communities (11 replicates) and negative controls containing a template of molecular grade water (11 replicates) were also processed and sequenced in parallel with the samples. Summary statistics for mock communities and negative controls are available in Table S3.

### Sequence processing

Demultiplexed sequences were trimmed with cutadapt (v. 2.10; Martin, 2011) then filtered and processed using the R (v. 4.2.1) package DADA2 (v. 1.26.0; Callahan et al., 2016). Reads were trimmed to 216 (forward reads) or 206 (reverse reads) base pairs, with those containing more than 2 expected sequencing errors removed. Amplicon sequence variants (ASVs) were then inferred from filtered reads. Paired reads were merged and length-filtered to remove sequences <320 or >380 bp, and bimera sequences were removed. Finally, SEPP ( v.4.5.1; Mirarab et al., 2011) was used to align the *nifH* sequences to the reference *nifH* alignment provided in the R package PPIT (Phylogenetic Placement for Inferring Taxonomy; v. 1.2.0; Kapili & Dekas, 2021).

From SEPP output files, PPIT was used to infer *nifH* taxonomy for each ASV. If the nearest reference to a query ASV was a *nifH*-like sequence, the query was also flagged as *nifH*-like. (*nifH*-like sequences were removed for all subsequent diazotrophic community analyses.) PPIT avoided inferring taxonomy and labeled an ASV “Unassigned” if the ASV did not share sufficient pairwise identity with references in the phylogenetic neighborhood, or if nearby reference sequences did not have consistent taxonomic classifications, a sign of horizontal gene transfer. PPIT therefore has a lower rate of identification compared to other methods, but higher accuracy (Kapili & Dekas, 2021).

### qPCR analysis

Quantitative PCR (qPCR) was performed with a StepOne Plus Real-Time PCR system (Applied Biosystems, Waltham, MA, USA) on the DNA and cDNA extracted from seep sediments. qPCR reactions had a total volume of 20 µL and consisted of 10 µL Takara TB Green Premix Ex Taq II (2x), 0.4 µL ROX dye, 0.8 µL of forward and 0.8 µL of reverse primers (10 µM concentration), 6 µL DNase-free water, and 2 µL DNA or cDNA template. The thermal cycling conditions followed the optimized protocol of Miyazaki et al., 2009, and were as follows: 95 °C for 120 s; 40 cycles of 96 °C for 30 s, 55 °C for 30 s, and 72 °C for 60 s; 95 °C for 15 s; 60 °C for 60 s, and 95 °C for 15s. Primers for qPCR were mehtaFw-28/mehtaRv-416 (Mehta et al., 2003) – the same used for *nifH* amplicon sequencing (and for qPCR in Dang et al., 2009, 2013; Miyazaki et al., 2009). Each sample was measured in triplicate. Concentrations per µL DNA/cDNA extract were converted to environmental concentrations copies per gram dry sediment using porosity values for each sample determined via weight loss after dehydration (and assuming a sediment density of 2.65 g cm^-3^). Porosity data reported for USAM samples in Semler et al. (2022) and for MB samples in Semler & Dekas (2024, preprint).

Quantification of *nifH* was based on a dilution series of *nifH* gene fragments synthesized from Twist Biosciences (South San Francisco, CA, USA) at a known concentration which was verified by Qubit. The gene fragments were obtained from the *nifH* sequence (KR020479.1) of a clone (CH4-cDNA-nifH-H5) originally discovered in the Costa Rican mud volcano Mound 12 (Dekas et al., 2016), and were diluted to concentrations ranging from 5 x 10^6^ to 0.5 *nifH* copies/µL. The limit of detection varied slightly between qPCR runs, with a mean of 1.30 x 10^2^ *nifH* copies/µL. Based on necessary sample dilutions (to conserve sample volume) and the DNA/RNA extraction process, this corresponded to a mean *environmental* detection limit of 3.12 x 10^5^ copies per gram dry sediment.

Gene and transcript concentrations were corrected for the presence of *nifH*-like genes by multiplying the raw copy number by the proportion of *nifH* reads that were bona-fide *nifH* sequences in that sample (as determined by the amplicon sequencing analysis using the same primer set). Corrected numbers are therefore lower than the raw values. When amplicon data was not available to correct raw qPCR values, abundances are not reported.

### Sequence analysis and statistical methods

Non-metric multidimensional scaling (NMDS) and hierarchical clustering of diazotrophic communities was carried out based on the weighted UniFrac distance metric (Lozupone et al., 2007) using the R package “vegan” (v. 2.5-7; Oksanen et al., 2020). Analysis of similarity (ANOSIM) was used to determine the significance of microbial community differences between groups of samples, also based on the weighted UniFrac distance metric. The statistical significance of all linear models was evaluated with a Bonferroni correction for multiple hypotheses testing.

## Results

### Nitrogen geochemical environment of cold seeps

Concentrations of nitrogen species (NH_4_ ^+^, NO_2_ ^-^, and NO_3_ ^-^) were measured in MB pushcores collected from seep and background sediments. Concentrations of NH_4_ ^+^, NO_2_ ^-^, and NO_3_ ^-^ from USAM sites were previously reported in Semler et al., 2022 and are included here for comparison. Overall, nitrogen species were higher in seep sediment relative to background sediment at MB sites, consistent with the trend found previously at Hydrate Ridge seeps (Dekas et al., 2018)) but lower in seeps relative to background sediments at USAM sites. Within and outside seeps, all nitrogen species were more elevated at MB sites than at USAM sites (Fig. 1). Ammonium typically decreased with depth in seep sediments and was particularly enriched at site MB-CF; concentrations reached 250 µM in the shallowest sediment depths there. Concentrations of nitrate also decreased with sediment depth within the seep boundaries, and was particularly high at MB-CF (Fig. 1). In background sediments, concentrations of all nitrogen species typically increased with depth, in contrast to the trend within seeps.

**Fig. 1:**
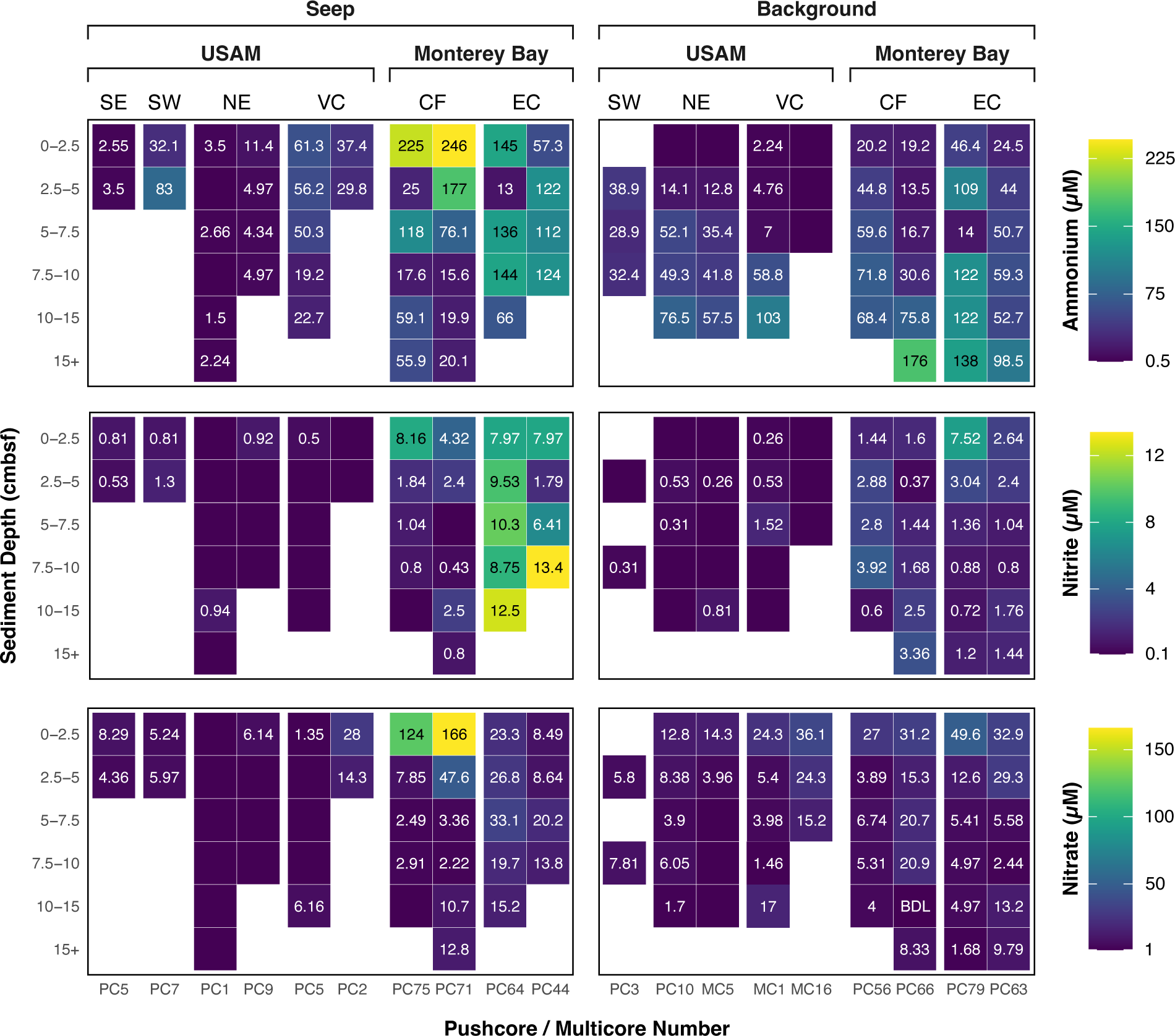
Ammonium, nitrite, and nitrate concentrations with sediment depth in all cores with geochemical data from inside and outside the six methane seep sites: Shallop Canyon East (SE), Shallop Canyon West (SW), New England (NE), Veatch Canyon (VC), Clam Field (CF), and Extrovert Cliff (EC). (Data from USAM locations previously reported in Semler et al., 2022.) White regions indicate no measurement was made. Blank values in filled boxes indicate a value below the detection limit of the assay. (Sediment horizon labels at USAM sites were altered due to space restrictions and for effective comparison. USAM sediment horizons were: 0-3, 3-6, 6-9, 9-12, 12-15, and 15+ cmbsf.)

### Detection of nifH gene and transcript sequences and nifH-like sequence removal

We successfully amplified and recovered *nifH* and *nifH*-like gene sequences from 97% of collected sediment samples (106 of 109), and transcript sequences from 22% of samples (24 of 109). In total, we generated 7,674,265 reads and inferred 36,803 unique amplicon sequence variants (ASVs) across all seep and background sediments. Using PPIT, we identified 70.8% of recovered ASVs as *nifH*-like sequences, which together accounted for 44.9% of total reads, leaving 10,734 bona fide *nifH* ASVs and 4,231,806 total reads after removal. In background sediments, 78.6% of ASVs and 86.9% of reads were *nifH*-like, while in seep sediments, a smaller percentage (57.8% of ASVs and 26.3% of reads) were *nifH*-like (Fig. 2). However, the percentage of *nifH*-like sequences in seep sediments was highly region-dependent. Only 41.1% of ASVs and 9.1% of reads in USAM seep sediments were identified as *nifH*-like, while 74.0% of ASVs and 72.2% of reads in MB seep sediments were *nifH*-like (Fig. 2). *nifH*-like sequences were not included in further analyses, and we use “*nifH*” to refer to bona fide *nifH* sequences only—those within *nifH* phylogenetic groups I, II, III, VII, and *Methanosarcina*-like (MSL)— herein (Fig. S2A-B). A recent report suggests that some *nifH* group IV (IV-A) sequences are also functional in *nifH* based on conserved protein residues (Dong et al., 2022), however, we conservatively exclude them from our classification of bona fide *nifH* sequences here.

**Fig. 2:**
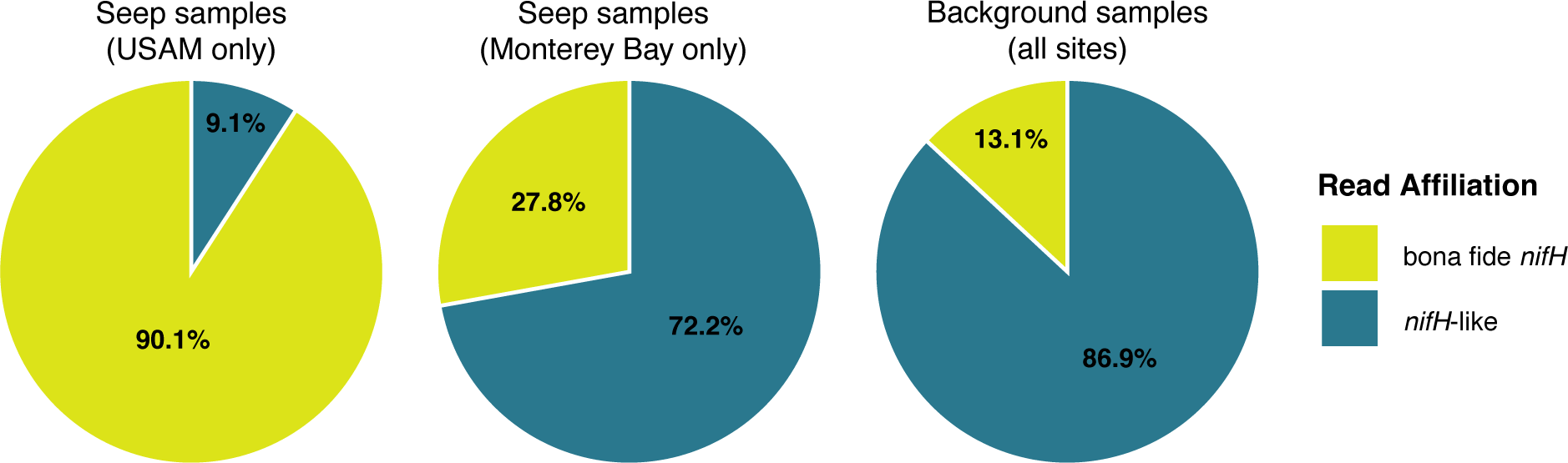
Percentage of reads generated with the mehtaFw-28/mehtaRv-416 (Mehta et al., 2003) primers that were affiliated with bona fide *nifH* (falling into *nifH* groups I, II, III, VII, or Methanoscarcina-like) vs. *nifH*-like sequences in USAM seep samples, in Monterey Bay seep samples, and in background samples.

### Distribution and diversity of nifH genes

The bona fide *nifH* gene ASVs recovered from within seep sediments (n = 5,799) spanned *nifH* groups I, II, III, VII, and the MSL group (Fig. S3A). We were able to assign taxonomy to 69.2% of these ASVs, which accounted for 89.3% of the *nifH* reads. These ASVs were affiliated with 17 unique phyla and 30 unique classes (Table S4), and varied widely by seep site and sediment depth. At USAM seeps, the most relatively abundant taxa were members of the predominantly anaerobic methane-oxidizing Methanomicrobia (Fig. 3A), *Candidatus* Methanoliparia (previously known as the GoM-Arc2 clade; Orcutt et al., 2010), Clostridia, and Deltaproteobacteria (now Desulfobacterota). *Ca.* Methanoliparia was present at sites USAM-NE and USAM-VC only. *nifH* sequences in MB were primarily affiliated with the Deltaproteobacteria – in particular, representatives of the orders Desulfuromonadales (mainly site MB-CF) and Desulfobacterales (both MB-CF and site MB-EC). Low abundance phyla present across all seep sites included members of the Kiritimatiellaeota, Lentisphaerae (now Lentisphaerota), Nitrospirae (now Nitrospirota), and Planctomycetes (now Planctomycetota) (Table S4).

**Fig. 3:**
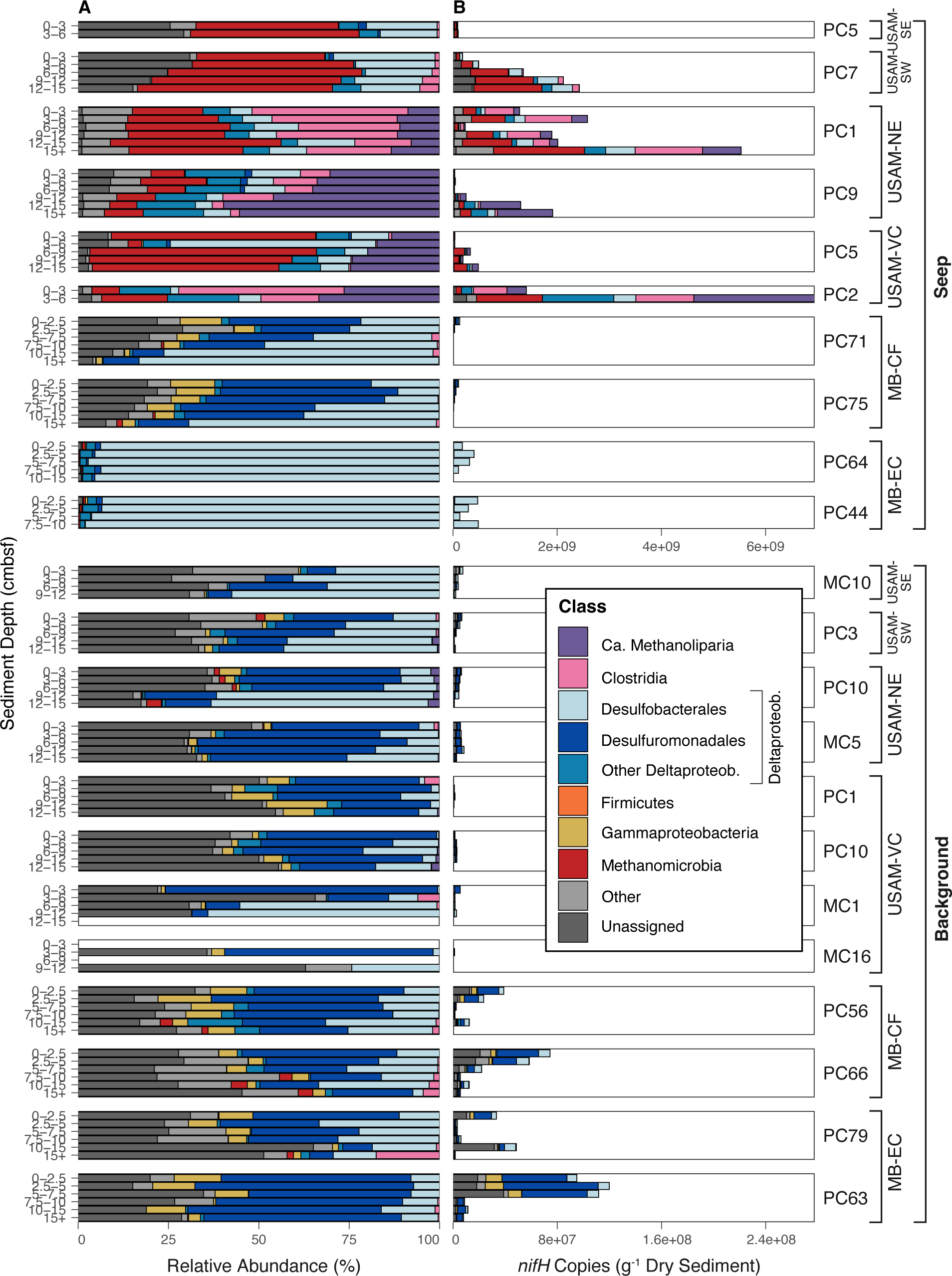
Relative abundance (A) and total abundance (B) of putative diazotrophs after removing *nifH*-like sequences, aggregated at the class or order (for Deltaproteobacteria) rank. Boxes surround each individual core and are labeled by pushcore/multicore number. Sediment depth increases from top to bottom in each core. Note: for total abundances in panel B, the x-axis is different for seep samples than for background samples. Other includes unclassified Deltaproteobacteria, Desulfovibrionales, Syntrophobacterales, and Myxococcales.

We recovered 5,578 *nifH* ASVs from the background sediments, also spanning *nifH* groups I, II, III, VII, and the MSL group (Fig. S3B). We were able to taxonomically assign 52.4% of these ASVs, comprising 65.5% of the background *nifH* reads. In these sediments from all sites, the identities of the NCOs and the patterns of their relative abundance and distribution broadly matched those at the MB seeps, particularly MB-CF seep (Fig. 3A). The NCOs spanned 18 unique phyla and 27 unique classes (Table S4), with assigned ASVs predominantly affiliated with the Deltaproteobacteria – particularly the Desulfuromonadales and the Desulfobacterales. In USAM-VC, MB-CF, and MB-EC background sediments, a small percentage of *nifH* sequences (∼5%) were affiliated with the Gammaproteobacteria (specifically the family Chromatiaceae, a group of nitrogen-fixing purple sulfur bacteria, which contain species that are capable of non-phototrophic growth (Imhoff, 2014). Low abundance phyla present across background sediments from all sites included members of the Acidobacteria (now Acidobacteriota), Bacteroidetes (now Bacteroidota), Chlorobi (now Chlorobiota), Kiritimatiellaeota, Lentisphaerae, Nitrospirae, and Planctomycetes (Table S4).

A non-metric multidimensional scaling (NMDS) ordination (Fig. S4) of the *nifH* sequences showed that the community of putative diazotrophs was primarily differentiated by site, which was found to be statistically significant by an ANOSIM test (R = 0.293, P = 0.001). Samples were also significantly differentiated by sediment type (seep or background; ANOSIM: R = 0.135; P = 0.001).Three general clusters of seep samples were apparent: one encompassing samples from USAM-SE and USAM-SW, one encompassing USAM-NE and USAM-VC, and one encompassing MB-CF and MB-EC (Fig. S4; Fig. S5).

### Detection of nifH genes from Euryarchaeota and Deltaproteobacteria

ANME-2 archaea represented a very large percentage of the identified NCOs within the cold seeps, comprising 30% of all *nifH* genes. However, their prevalance varied dramatically between sites. At USAM seeps, ANME-2 comprised 36% of the identified NCOs on average, while at MB seeps, they comprised only 0.3%. In background sediments, ANME-2 also comprised only 0.3% of NCOs. The ASVs affiliated with ANME-2 were all extremely similar to one another, and the majority clustered together within the MSL-like *nifH* group (Fig. 4). *nifH*-like sequences affiliated with ANME-1 were detected within seep samples, consistent with previous work that classified ANME-1 sequences as part of *nifH* group IV, non-functional in nitrogen fixation (Meyerdierks et al., 2010), but potentially functional in the coenzyme F430 biosynthetic pathway (Ghebreamlak & Mansoorabadi, 2020).

**Fig. 4:**
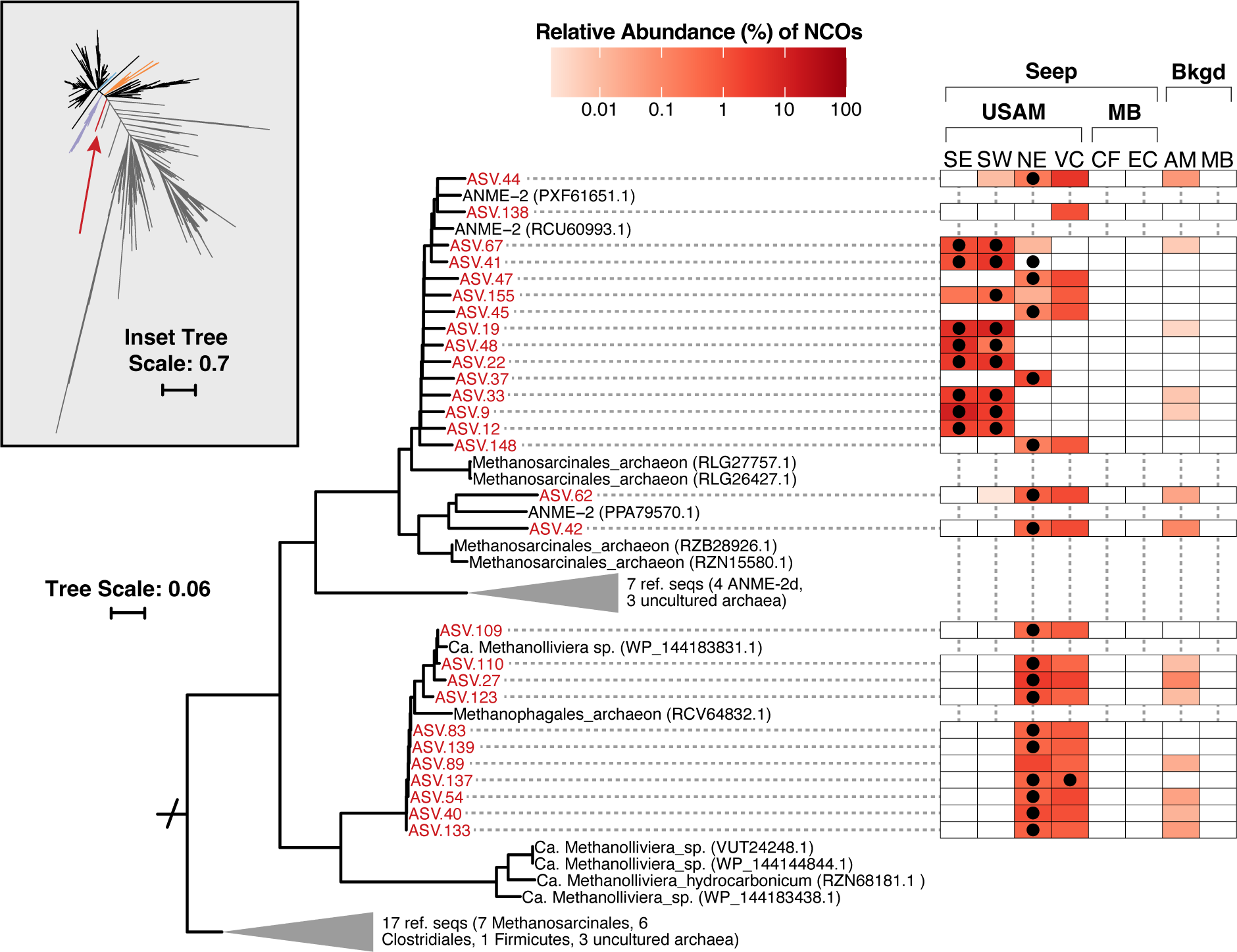
SEPP-derived phylogenetic placement of *nifH* ASVs assigned to the ANME-2 or *Ca.* Methanoliparia (n = 42 ASVs), and their relative abundance (DNA) among all *nifH*-containing organisms (NCOs) at the six seep sites and in the two background sediment regions. (All background sediments collapsed into region regardless of site.) Only ASVs within the 100 most abundant were included in the phylogenetic tree for clarity. Taxonomic assignments were based on the package PPIT; ASVs assigned to ANME-2 or *Ca.* Methanoliparia are labeled in red. Black dots represent ASVs present in *nifH* transcripts. Inset shows the full *nifH* reference tree from the PPIT package and Supplementary Fig. S2A, with a red arrow pointing to the clade containing the inferred ANME or *Ca.* Methanoliparia ASVs. Scale bars show the expected number of nucleotide substitutions per site. Number of samples collapsed: Shallop Canyon East (SE; n = 2 gene samples, n = 2 transcript samples), Shallop Canyon West (SW; n = 5 gene, n = 5 transcript), New England (NE; n = 12 gene, n = 8 transcript), Veatch Canyon (VC; n = 7 gene, n = 2 transcript), Clam Field (CF; n = 12 gene, n = 0 transcript), Extrovert Cliff (EC; n = 9 gene, n = 6 transcript), USAM background (AM; n = 35 gene, n = 1 transcript), Monterey Bay background (MB; n = 24 gene, n = 0 transcript).

ASVs affiliated with *Ca.* Methanoliparia were also discovered at high relative abundances, with an average of 26% at USAM-NE and USAM-VC seeps, though they only comprised 0.03% of other seep sites on average (Fig. 4). USAM-NE and USAM-VC were the two deepest seep locations overall, at roughly 1,150 and 1,500 mbsl, respectively, and the abundance of *Ca.* Methanoliparia *nifH* genes increased with sediment depth at both sites (Fig. 3A).

Within the Deltaproteobacteria, sequences from two orders – Desulfobacterales and Desulfuromonadales – were the primary diazotrophic groups at seep and background sites (Fig. 3A). ASVs classified as Desulfobacterales were split between 2 separate branches of the *nifH* reference tree (Fig. 5A-B). The first branch – the branch containing many Group III, VII, and MSL-like *nifH* sequences (Fig. S2A) – was primarily found at MB seeps (Fig. 5A), and to a lesser extent at USAM-NE and USAM-VC seeps. Three of the ASVs (ASV.20 at MB-CF and ASV.3 and ASV.17 at MB-EC) each comprised >25% of the *nifH* sequences found at their respective sites. These ASVs were all closely related to *Desulfoglaeba alkanexedens*, an organism capable of nitrogen fixation that also oxidizes n-chain alkanes in oily environments (Davidova et al., 2006). ASVs related to *D. alkanexedens* were not found at USAM seeps but were found to a lesser degree (<1% relative abundance) in MB background sediments. The second group of Desulfobacterales was located on a branch of the *nifH* tree containing many Group II *nifH* sequences (Fig. S2A). ASVs were placed close to unspecified Desulfobacterales and Desulfobacteraceae reference sequences and were found at the highest abundances at the USAM-SE and USAM-SW seep sites (Fig. 5B).

**Fig. 5:**
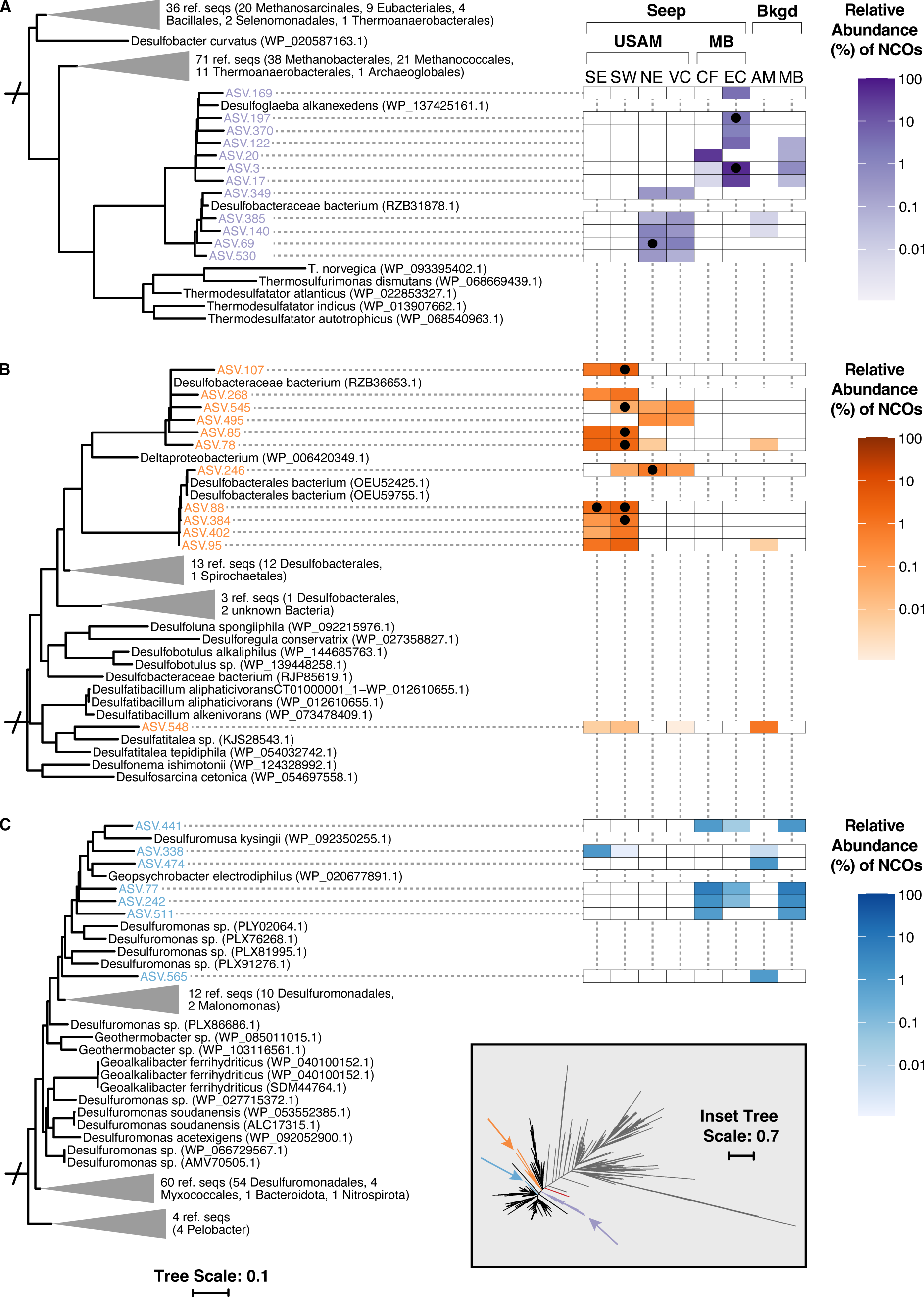
SEPP-derived phylogenetic placement of *nifH* ASVs assigned to the Desulfobacterales (n = 24 ASVs) or Desulfuromonadales (n = 7 ASVs), and their relative abundance (DNA) among all *nifH*-containing organisms (NCOs) at the six seep sites and in the two background sediment regions. (All background sediments collapsed into region regardless of site.) Only ASVs within the 500 most abundant were included in the phylogenetic tree for clarity. Taxonomic assignments were based on the package PPIT; ASVs assigned to Desulfobacterales are labeled in either purple (A) or orange (B), depending on the reference tree source branch, and ASVs assigned to Desulfuromonadales are labeled in blue (C). Black dots represent ASVs present in *nifH* transcripts. Inset shows the full *nifH* reference tree from the PPIT package and Supplementary Fig. S2A, with purple and orange arrows pointing to the clades containing the inferred Desulfobacterales ASVs and blue arrows pointing toward inferred Desulfuromonadales ASVs. Scale bar shows the expected number of nucleotide substitutions per site. Number of samples collapsed: Shallop Canyon East (SE; n = 2 gene samples, n = 2 transcript samples), Shallop Canyon West (SW; n = 5 gene, n = 5 transcript), New England (NE; n = 12 gene, n = 8 transcript), Veatch Canyon (VC; n = 7 gene, n = 2 transcript), Clam Field (CF; n = 12 gene, n = 0 transcript), Extrovert Cliff (EC; n = 9 gene, n = 6 transcript), USAM background (AM; n = 35 gene, n = 1 transcript), Monterey Bay background (MB; n = 24 gene, n = 0 transcript).

Desulfuromonadales ASVs alone comprised 5.4% of all *nifH* genes across our dataset and were particularly abundant in background sediments (30%), as well as in the seep sediments of site MB-CF (27%). Regardless of source sediment, most Desulfuromonadales ASVs clustered into a single clade (Fig. 5C), which also included reference sequences from *Desulfuromusa kysingii* and *Geopsychrobacter electrodiphilus*. These organisms are obligate anaerobes known to oxidize fermentation products (especially acetate, propionate, amino acids, and aromatic compounds) by reducing iron oxides or elemental sulfur (Holmes et al., 2004; Liesack & Finster, 1994; Vandieken et al., 2006).

### Distribution and diversity of nifH transcripts

*nifH* transcripts were detected in 22% of the samples in our dataset, comprising 948 ASVs. Transcripts were only successfully amplified in seep samples, with a single exception in a background sample (site USAM-NE, PC10, 12-15 cmbsf). The transcripts spanned *nifH* groups X-X. Their inferred hosts were far less taxonomically diverse than the hosts of all *nifH* genes, spanning only 3 phyla, the Euryarchaeota, Firmicutes (now Bacillota), and Proteobacteria [now Pseudomonadota, (Table S5)], despite a higher percentage of taxonomic assignment of transcripts (97.7% of total transcript reads assigned) than of genes (86.5% of total gene reads assigned) (Fig. 4). Transcripts were recovered from all USAM seeps, and were largely affiliated with ANME-2, though transcripts associated with *Ca.* Methanoliparia, Clostridia, and Deltaproteobacteria—the latter unclassified beyond the class rank—were also detected (Fig. 4; Fig. 5; Table S5). At site MB-EC, the majority of transcripts amplified were affiliated with Desulfobacterales (Fig. 5A-B), with some also from Desulfuromonadales (Fig. 5C). A lower percentage of transcripts were assigned taxonomy at MB than at USAM (79.7% versus 98.2%), potentially reflecting the presence of diazotrophs more poorly represented in databases. No *nifH* transcripts were amplified at site MB-CF.

### Quantification of nifH genes and transcripts

We quantified *nifH* via qPCR to investigate the abundance (DNA) and potential activity (cDNA) of putative diazotrophs across sites, sediment depths, and environmental variables. Abundances were corrected for the presence of *nifH*-like genes and transcripts, as determined with the sequencing data (see Methods), decreasing the raw values by 0-99.9% per sample (Fig. S6). We detected *nifH* genes and transcripts in 96% and 35% of samples, respectively; greater than that with the amplicon approach. Where detected, corrected concentrations of *nifH* genes and transcripts within seeps ranged from 4.4 x 10^6^ to 6.2 x 10^9^ (Fig. 3B; Fig S7) and 1.4 x 10^5^ to 3.8 x 10^6^ copies per gram dry sediment (Fig. 6B; Fig. S7), respectively. Abundances of *nifH* at USAM seeps (particularly at site USAM-NE and site USAM-VC) were the highest (1.4 x 10^9^ and 1.3 x 10^9^ on average, respectively) and increased with sediment depth. At seeps from site MB-CF, abundances of *nifH* actually decreased with depth, while at MB-EC, *nifH* abundance had no discernible depth trend. We detected transcripts via qPCR at all seep sites except MB-CF, and found the highest concentrations at USAM-SW (4.1 x 10^6^ at site).

**Fig. 6:**
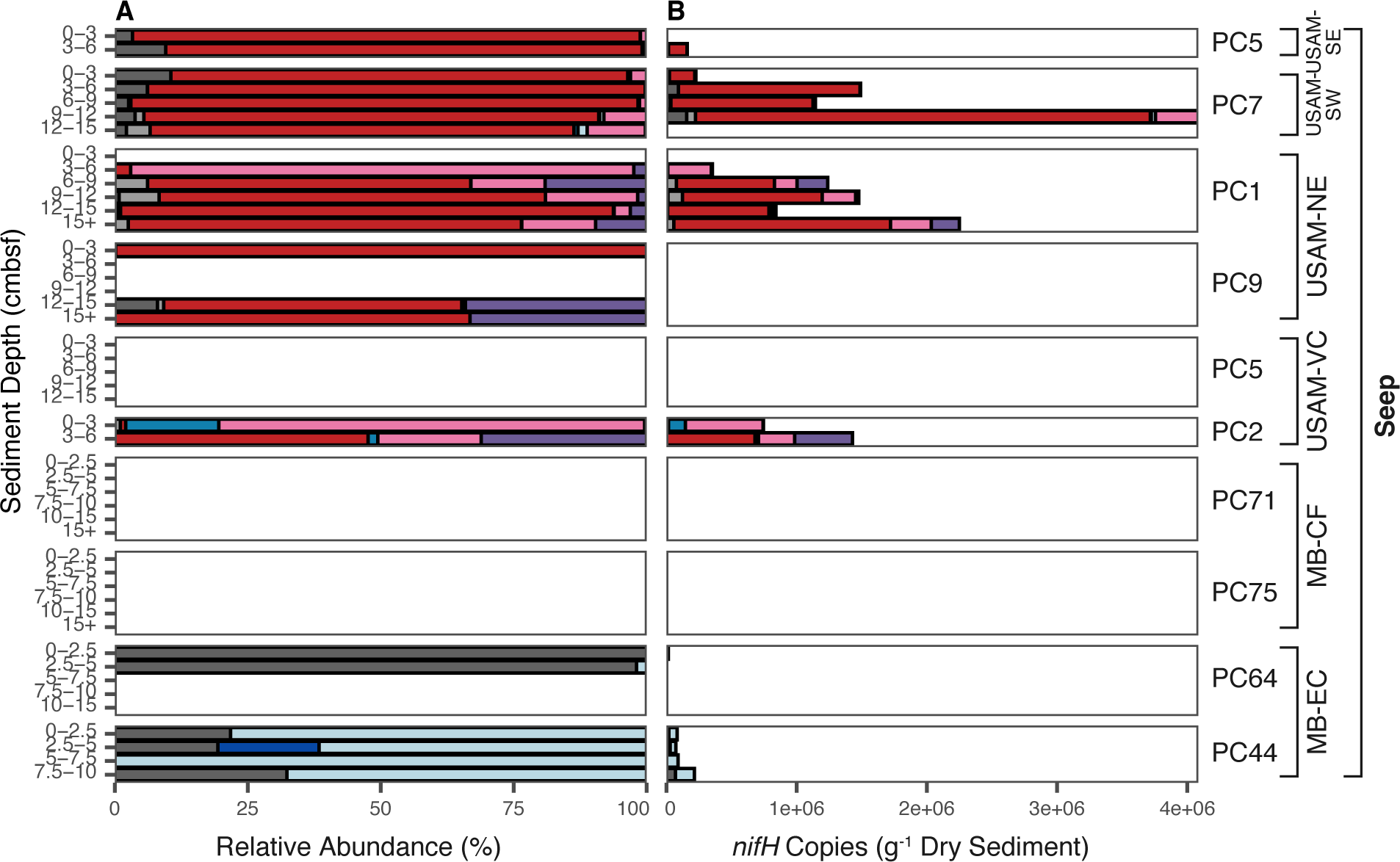
Relative abundance (A) and total abundance (B) of *nifH* transcripts after removing *nifH*-like sequences, aggregated at the class or order (for Deltaproteobacteria) rank. Boxes surround each individual core and are labeled by pushcore/multicore number. Sediment depth increases from top to bottom in each core. Note: for total abundances in panel B, the x-axis is different for seep samples than for background samples. Legend colors as in Fig. 3.

In background sediments, concentrations of *nifH* genes ranged from 9.7 x 10^5^ to 9.6 x 10^9^ copies per gram dry sediment – four orders of magnitude. Copies of *nifH* decreased with sediment depth, and were found to be most abundant in MB background sediments. *nifH* gene and transcript concentrations were generally lower in background sediments than at cold seeps by 1 to 2 orders of magnitude, and up to 3. Background sediments did not contain detectable concentrations of *nifH* transcripts, with the exception of 11 samples at site MB-CF, which were detected by qPCR but did not have corresponding amplicon sequencing data to correct values, and are therefore not reported. (Fig. S7).

### Correlations between nifH gene abundances and biogeochemistry

To determine potential drivers of diazotroph distribution at cold seeps, the abundance of *nifH* genes in each seep sample was compared to the concentrations of a variety of geochemical species (CH_4_, S^2-^, NH_4_ ^+^, NO_2_ ^-^, and NO_3_ ^-^). None of these species were significantly correlated with *nifH* gene abundance (Fig. S8). However, due to the inhibition of nitrogen fixation that has been observed above ammonium concentrations of 25 µM in both pure cultures (Kessler et al., 2001) and in incubations of diverse marine sediment samples (Dekas et al., 2018), we also investigated the relationship between *nifH* gene abundance and ammonium concentration separately for “low” (< 25 µM) and “high” (> 25 µM) ammonium samples. As we predicted, this separation revealed a negative correlation between *nifH* genes and ammonium in seep samples below the threshold of ammonium inhibition (r^2^ = 0.525, p = 6.0 x 10^-3^), and resulted in no significant trend in seep samples above it (Fig. 7). In background sediments, there was no significant trend between ammonium and *nifH* genes – neither in high ammonium samples nor in low ammonium samples (p > 0.05) (Fig. 7). The significant relationship between *nifH* gene abundance and ammonium concentration at seeps was not observed before raw gene abundances were corrected to remove *nifH*-like sequences, highlighting the importance of filtering the *nifH*-like sequences.

**Fig. 7:**
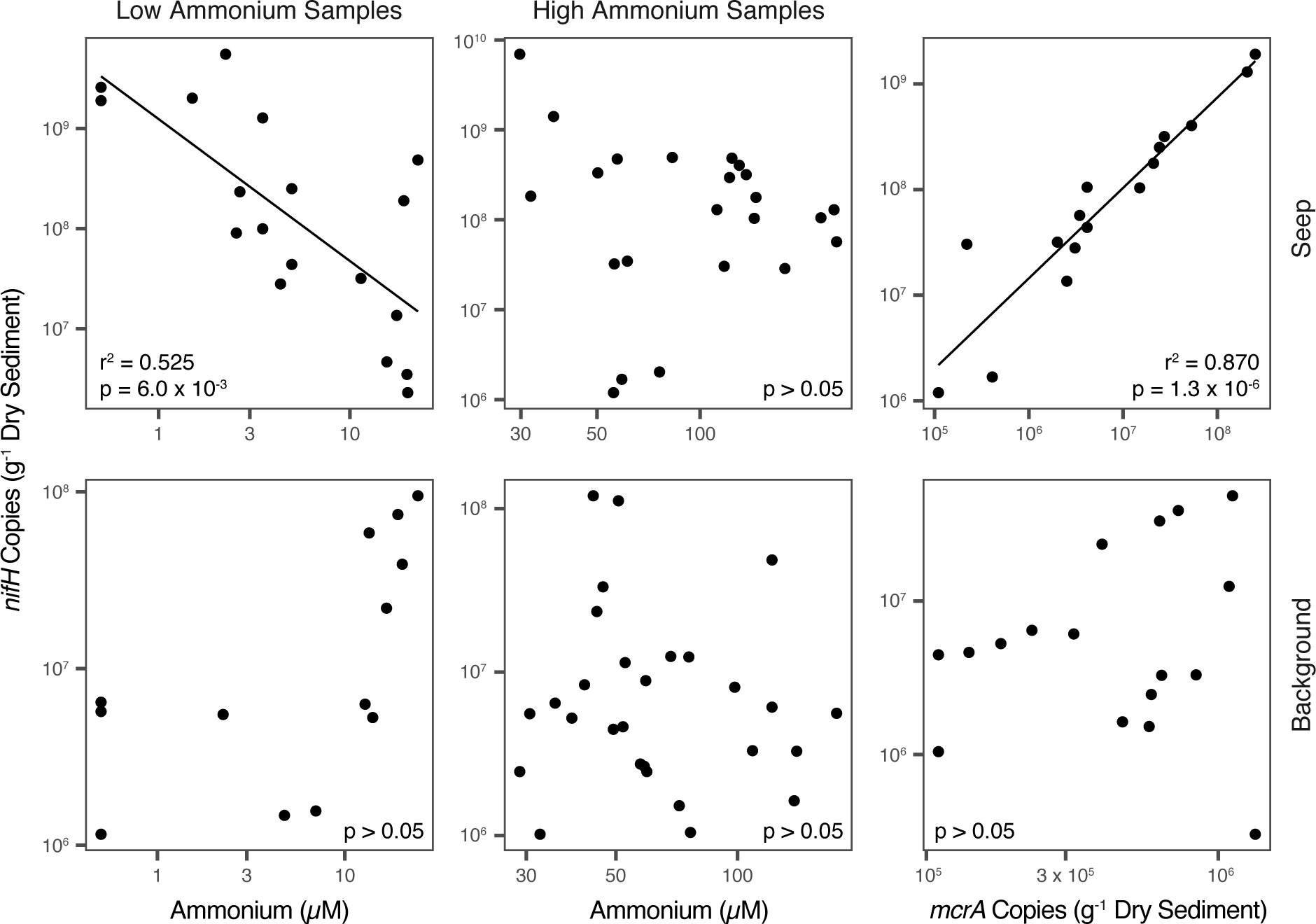
Corrected *nifH* gene abundances in seep or background samples compared to ammonium concentrations in both low ammonium (< 25 µM) and high ammonium (> 25 µM) samples, and to *mcrA* gene abundances. Significant relationships (p-value < 0.05) shown with solid trendlines; those with no significance (p-value > 0.05) shown with no trendlines.

The abundance of *nifH* genes in seep samples was also highly correlated with the abundance of *mcrA* genes previously reported in Semler & Dekas (2024, preprint) (Fig. 7; r^2^ = 0.870, p = 1.3 x 10^-6^). In background sediments, a correlation between these two genes was not observed. The abundance of *nifH* transcripts in seep and background samples was not significantly correlated with any of the examined variables, perhaps due to the low number of transcript samples with paired geochemistry.

## Discussion

### Major diazotrophic assemblages at marine cold seeps

Within sediment from six seeps in two oceans, we recovered 5,799 unique bona fide *nifH* sequences spanning nifH groups I, II, II, VII, and MSL. These sequences were affiliated with 17 different bacterial and archaeal phyla, significantly exanding the known diversity of NCOs at cold seeps. Between sites, there were substantial shifts in taxonomic identity of the NCOs (Fig. 3A; Fig. 6A). Seeps in our dataset contained one of three distinct diazotrophic assemblages (Fig. S5). The first assemblage, which was present at USAM-SE and USAM-SW seep sites, was comprised of ANME-2 archaea and sulfate reducing bacteria (SRB) of the Desulfobacteraceae (from the order Desulfobacterales) and comprised 53% and 61%, respectively, of the *nifH* genes at those sites. Desulfobacteraceae include the Seep-SRB1a clade, which are obligate syntrophic partners of the ANME-2 (Kleindienst et al., 2014; Knittel et al., 2003; Schreiber et al., 2010) and were previously found at high abundances in 16S rRNA gene surveys of these two sites (Semler et al., 2022). Sequences from this diazotrophic assemblage have dominated previous *nifH* clone libraries from seeps (Dang et al., 2013; Dekas et al., 2016; Miyazaki et al., 2009), and have comprised a large percentage of the *nifH* genes recovered in seep MAGs (Dong et al., 2022). ANME and Desulfobacteraceae groups also dominated *nifH* transcripts from USAM-SE and USAM-SW, comprising 94% and 89%, respectively, of *nifH* transcripts (Fig. 6). At USAM-SE and USAM-SW, the syntrophic consortium between ANME and their SRB partners are likely the most abundant and active diazotrophs, despite the high energetic costs of diazotrophy for organisms performing the anaerobic oxidation of methane.

A second major diazotrophic assemblage at our seeps also contained the ANME-2/SRB consortia, with the addition of archaea from the candidate class *Ca.* Methanoliparia within the Euryarchaeota (Fig. S5). Based on its genomic composition, *Ca.* Methanoliparia has been implicated in the methanogenic degradation of short-chain alkanes (Laso-Pérez et al., 2019), but has not been previously recognized as a potential diazotroph despite encoding *nifH*. Culture-based experiments also confirmed that this group is capable of breaking down long-chain and aeromatic hydrocarbons into carbon dioxide and methane (Zhou et al., 2022). *Ca.* Methanoliparia accounted for 26% of the *nifH* genes at USAM-NE and USAM-VC seeps (with ANME-2 and Desulfobacteraceae comprising an additional 37%), indicating that these two sites may have input of non-methane hydrocarbons, particularly at greater sediment depths where *Ca.* Methanoliparia comprises a larger percentage of NCOs. *Ca.* Methanoliparia also comprised more than 10% of *nifH* transcripts from USAM-NE and USAM-VC seeps on average (Fig. 6), indicating that these organisms not only encode *nifH*, but also are potentially active in diazotrophy.

The third and final major diazotrophic assemblage at our seep sites was found at site MB-CF and site MB-EC, and predominantly contained putatively alkane-oxidizing sulfate reducers of the Desulfobacterales, as well as iron-reducing members of the Desulfuromonadales (Fig. S5). Transcripts of two of these Desulfobacterales ASVs (though none of the Desulfuromonadales ASVs) were also recovered from MB-EC (Fig. 5). Oil and other hydrocarbons have been found at MB seeps (Lorenson et al., 2002; Orange et al., 1999; Stakes et al., 1999), and seep fluids are thought to be sourced from the underlying Monterey Formation, which contains abundant hydrocarbons. In addition, ANME archaea were found to be nearly absent in 16S rRNA sequencing surveys of these seep sites (Semler & Dekas, 2024, preprint), making their absence from *nifH* sequencing surveys in MB sediments predictable, if unusual.

### *Phylogenetically diverse, low abundance NCOs* and the true diversity of diazotrophs at seeps

Our dataset reveals a great diversity of *nifH* sequences and affiliated hosts, far beyond what has been observed at seeps previously with clone libraries or metagenomics, and also beyond surveys of high throughput amplicon sequencing in the marine watercolumn (Morando et al., 2024, preprint). Beyond the highly abundant NCOs in the Euryarchaeota and Proteobacteria, described above, we detected *nifH* sequences affiliated with 15 other bacterial and archaeal phyla at lower concentrations (Table S4; Table S5), several of which were present at all 6 seep sites. Transcripts of the majority of these phyla were not detected here, but representatives of some of these same phyla were identified as active diazotrophs in deep-sea sediment outside of seeps in a previous study (Kapili et al., 2020). Since our approach to assigning taxonomy to *nifH* sequences requires a reference database, we are unable to infer *nifH* in organisms without a previously published genome containing it. Sequences from novel diazotrophs, along with those in areas of the *nifH* phylogenetic tree with inconsistent host identity, fall into the “Unassigned” category of our analysis. This represents about 10.7% or 34.5% of all DNA reads in seep and background sediments, respectively, and 2.3% of all cDNA reads. The true diversity of *nifH* sequences and NCOs at our sites therefore extends beyond the groups reported here, and suggests marine sediments may be a rich source of novel diazotrophs. Total microbial community composition at seeps is marked by high levels of local diversification due to environmental selection (Ruff et al., 2015; Semler et al., 2022) but a high degree of functional similarity across sites, perhaps explaining the phylogenetic heterogeneity of putative nitrogen fixers.

Conversely, the presence of “orphan” *nifH* sequences could inflate the diversity of diazotrophs in our dataset. These are *nifH* sequences which occur in genomes in the absence of the other genes necessary to encode nitrogenase (e.g., nifD, nifK), and are estimated to represent up to 20% of NCOs in some environments (Mise et al., 2021). While PPIT can successfully identify *nifH*-like sequences (Kapili & Dekas, 2021), orphan *nifH* sequences cannot be reliably identified by phylogeny, and would therefore be included in our analysis. However, orphan *nifH* sequences mainly belong to the class Clostridia, or to select methanogenic lineages – namely *Methanobrevibacter* and *Methanocaldococcus* (Mise et al., 2021), which represent a minority of the sequences recovered here (Fig. 3). Furthermore, we detected *nifH* transcripts of Clostridia at USAM seep sites (Fig. 6; Table S5), suggesting functionality in certain members of this group. While our results therefore likely largely reflect the diazotrophic community, a metagenomic analysis would be necessary to confirm the presence of all necessary *nif* genes for nitrogen fixation in individual taxa. In practice, this would not be possible, because metagenomic sequencing does not recover nearly the depth of *nifH* sequences as an amplicon approach [e.g., 10,734 *nifH* ASVs here versus 66 in Dong et al. (2022)]. These approaches are therefore complimentary, with amplicon analysis offering a broad perspective of potential diazotrophs, and metagenomics offering deeper insight into fewer organisms, as well as the ability to identify novel NCOs. Exploring the identity of the unknown NCOs detected here, as well as the nitrogen-fixing capacity of all of these NCOs—novel or not—will further refine the scale of diazotrophic diversity in marine sediments.

### Gene abundance of nifH within cold seeps

Our results stress that raw *nifH* copy numbers obtained through qPCR with the commonly used mehtaFw-28/mehtaRv-416 primers need to be regarded as maximum potential abundances, and likely overestimates, due to the inclusion of non-*nifH* targets (*nifH*-like sequences). In the case of background sediments, almost 90% of the reads identified by this widely used primer set were *nifH*-like sequences (Fig. 2), which would have led to an overestimate of *nifH* gene presence by nearly tenfold if left uncorrected. Similarly, in seep sediments, the presence of *nifH*-like sequences varied significantly, indicating that the type of environment does not reliably predict the level of *nifH*-like contamination. While MB-CF and MB-EC seeps appeared at first glance to contain *nifH* genes in similar abundance to USAM seep sites, abundances were much lower when the proportion of sequences affiliated with suspected homologs was removed from the dataset (Fig. S6). The mehtaFw-28/mehtaRv-416 primers are generally considered the best option to survey *nifH* in communities dominated by non-photosynthetic diazotrophs due to their robust performance and sequence inclusivity particularly with *nifH* group II and III sequences (Gaby & Buckley, 2012; Kapili et al., in prep). The undesired inclusion of *nifH*-like sequences is not a problem for sequencing studies, which can easily filter these out, but is more problematic for qPCR applications. Coupling qPCR and sequencing, as done here, can facilitiate a correction of the abundance data.

Corrected *nifH* gene concentrations ranged from 6.0 x 10^5^ to 1.3 x 10^9^ copies per gram wet sediment across our six cold seeps, which is equivalent to 1.2 x 10^6^ to 6.9 x 10^9^ copies per gram dry sediment (Fig. 7). This extends the range previously reported at seeps, with 10^8^ copies per gram wet sediment detected in Kumano Knoll (Miyazaki et al., 2009) and 10^6^ copies per gram wet sediment in various methanic South China Sea locations (Dang et al., 2013). These previous studies used the same qPCR primers but did not correct values to remove *nifH*-like sequences. The observed 1-3 order of magnitude increase between *nifH* gene concentrations in background vs. seep sediments here (Fig. 3, Fig. S6) was previously seen in comparisons of various South China Sea sites, but not reported across such small distances (e.g. 5-500m seep to background here, versus 10s to 100s of kilometers).

### Biogeochemical controls on nifH gene abundances

The wide (three orders of magnitude) range in *nifH* gene concentrations in seep sediments (Fig. 3; Fig. S6) – demonstrates that the capacity for diazotrophy at cold seeps is heterogeneous. In MB samples, the number of *nifH* genes was only slightly higher with hydrocarbon input (within the seep) than without (in nearby background sediment), while *nifH* gene abundance in USAM seep samples was much higher than in the surrounding sediment. Hydrocarbon concentration has been previously suggested as a potential control on *nifH* gene abundances (Dong et al., 2022) at seeps, based on the high density of diazotrophs often seen at high-flux seep systems. However, methane concentration was not significantly correlated with *nifH* gene abundance at our sites (Fig. S8). In fact, site MB-EC contained the highest methane concentrations (Supplementary Dataset S3.1), but one of the lowest concentrations of *nifH* genes (Fig. 3; Fig. S7). Given the highly significant correlation we observed between *nifH* and *mcrA* gene abundances (Fig. 7), however, it is possible that the previously reported link between *nifH* and methane actually reflects a link between *nifH* and the abundance of methanotrophs. Methanotroph abundance is dependent on a variety of factors, including cm-scale co-location of methane and sulfate, and intrigingly does not always correlate with methane concentrations at the site-level (Semler and Dekas, submittted).

Our data also suggest a strong negative correlation between *nifH* gene abundance and ammonium concentration (Fig. 7) at seeps. The seep pushcores with the highest densities of putative diazotrophs (namely PC1 from USAM-NE and PC2 from USAM-VC), were also the seeps where bioavailable nitrogen was the lowest (Fig. 1). The correlation between *nifH* gene abundance and ammonium concentration was only significant below 25 µM ammonium – the threshold for inhibition of diazotrophic activity (Dekas et al., 2018; Kessler et al., 2001). Above concentrations of 25 µM, ammonium did not affect the abundance of diazotrophs, as would be expected if high concentrations of ammonium removed the selective advantage of the ability to fix nitrogen. This might explain the lower concentrations of *nifH* genes in MB seep sediments, which are located in a submarine canyon close to terrestrial sources of nitrogen and along a coast with significant nutrient upwelling (Kudela & Dugdale, 1996; Pennington & Chavez, 2000), and therefore replete in ammonium.

The negative correlation we observed between *nifH* gene abundance and ammonium concentration in sediments is opposite the trend previously observed at seeps in the South China Sea (Dang et al., 2013), where *nifH* abundance (also quantified using the mehtaFw-28/mehtaRv-416 primer set) was instead positively correlated with porewater ammonium concentration. However, we did not observe a statistically significant negative correlation until *nifH*-like reads were removed from our analysis, demonstrating that this correction is necessary to reveal trends between geochemistry and the capacity for nitrogen fixation. This is particularly true when many *nifH* gene homologs, like those involved in coenzyme F430 biosynthesis, are known to be common in seep-dwelling methanogens and methanotrophs.

## Conclusion

In total, our analysis provides a spatially-resolved view of the diversity, abundance, and transcriptional activity of putative diazotrophs inside and outside marine cold seeps. Our findings reveal a wide phylogenetic diversity among NCOs at cold seep sites, but also indicate that most NCOs at seeps (including ANME-2/SRB consortia, *Ca.* Methanoliparia, and putatively alkane-oxidizing sulfate reducers of the Desulfobacterales) are likely directly involved in methane oxidation or the degradation of more complex hydrocarbons. Even at seeps such as those in MB that lack ANME archaea, there was a large overlap between hydrocarbon- and nitrogen-cycling. Additionally, our study reveals the significant impact of *nifH*-like sequence removal on our understanding of diazotrophic abundance and distribution; the anticorrelation between *nifH* gene abundance and ammonium concentration, in particular, was not observed before *nifH*-like sequences were eliminated. Moving forward, it will be imperative for future studies to account for contamination of *nifH*-like genes in *nifH* qPCR datasets, both in cold seep sediments, and elsewhere.

## Supporting information

Supplementary Datatset S1

## Acknowledgments

We thank the captain, crew, and science party of R/V *Atlantis* AT36 and R/V *Western Flyer* MMV19, as well as the pilots and engineers of HOV *Alvin* and ROV *Doc Ricketts*. We also thank all participants and mentors on the UNOLS Early Career Training Cruise AT36 for assistance in cruise planning and sample collection, and the members of the Dekas Geomicrobiology Lab for discussions and feedback – particularly Rebecca Salcedo for her metagenomics expertise. Funding was provided by Stanford University (to AS and AD) and NSF (OCE-1634297 to AD). The UNOLS Early Career Training Program was funded by NSF (OCE-1641453, OCE-1638805, OCE-1214335, OCE-1655587, and OCE-1649756) and ONR (N00014–15-1–2583).

## Data Availability

Nucleotide sequences from this study were deposited in the European Nucleotide Archive, project number PRJEB72160. Sample metadata is contained in Supplementary Dataset S1.

---

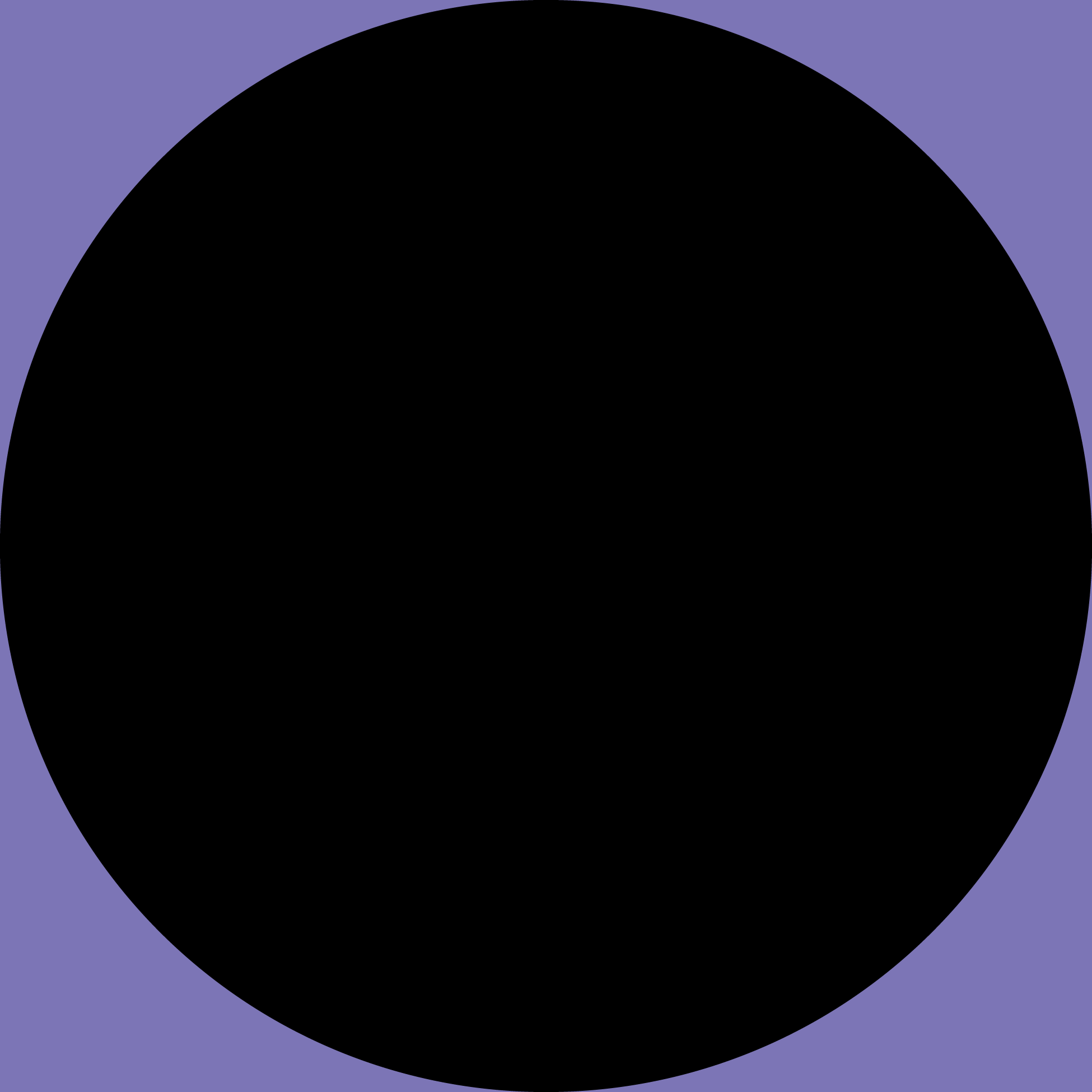

