## Supplementary Tables and Figures for "Abundance, identity, and potential diazotrophic activity of *nifH*-containing organisms at marine cold seeps"

Table S1: Sediment core information.

| Site Name | Core Type | Dive Number | Core Number | Latitude; Longitude | Water Depth (m) | Sediment Horizons Sampled (cmbsf) | Substrate Type |
|---|---|---|---|---|---|---|---|
| Shallop Canyon East | Seep | AD4832 | PC 5 | 39.995; -69.1287 | 366 | 0-3, 3-6 | Patchy microbial mat |
| Shallop Canyon East | Bkgd | N/A | MC 10 | 40.0103; -69.1433 | 335 | 0-3, 3-6, 6-9, 9-12 | Sand |
| Shallop Canyon West | Seep | AD4829 | PC 7 | 39.9692; -69.1933 | 395 | 0-3, 3-6, 6-9, 9-12, 12-15 | Patchy microbial mat |
| Shallop Canyon West | Bkgd | AD4829 | PC 3 | 39.9690; -69.1950 | 390 | 0-3, 3-6, 6-9, 9-12, 12-15 | Sand |
| New England | Seep | AD4834 | PC 1 | 39.9018; -69.2568 | 1150 | 0-3, 3-6, 6-9, 9-12, 12-15, 15+ | Patchy microbial mat |
| New England | Seep | AD4834 | PC 9 | 39.9030; -69.2520 | 1141 | 0-3, 3-6, 6-9, 9-12, 12-15, 15+ | Patchy microbial mat and mussels |
| New England | Bkgd | AD4834 | PC 10 | 39.9025; -69.2528 | 1130 | 0-3, 3-6, 6-9, 9-12, 12-15 | Sand |
| New England | Bkgd | N/A | MC 5 | 39.9019; -69.2638 | 1252 | 0-3, 3-6, 6-9, 9-12, 12-15 | Sand |
| Veatch Canyon | Seep | AD4828 | PC 5 | 39.8049; -69.5935 | 1420 | 0-3, 3-6, 6-9, 9-12, 12-15 | Patchy microbial mat |
| Veatch Canyon | Seep | AD4835 | PC 2 / PC 8 (paired) | 39.8103; -69.5901 | 1408 | 0-3, 3-6 | Thick mussel bed |
| Veatch Canyon | Bkgd | AD4828 | PC 1 | 39.8054; -69.5940 | 1407 | 0-3, 3-6, 6-9, 9-12, 12-15 | Sand |
| Veatch Canyon | Bkgd | AD4828 | PC 10 | 39.8066; -69.5926 | 1409 | 0-3, 3-6, 6-9, 9-12, 12-15 | Sand |
| Veatch Canyon | Bkgd | N/A | MC 1 | 39.8041; -69.5877 | 1496 | 0-3, 3-6, 6-9, 9-12, 12-15 | Sand |
| Veatch Canyon | Bkgd | N/A | MC 16 | 39.8068; -69.5985 | 1545 | 0-3, 3-6, 6-9, 9-12 | Sand |
| Clam Field | Seep | DR1139 | PC 75 | 36.7356; -122.0342 | 895 | 0-2.5, 2.5-5, 5-7.5, 7.5-10, 10-15, 15+ | Patchy microbial mat |
| Clam Field | Seep | DR1139 | PC 71 | 36.7356; -122.0342 | 895 | 0-2.5, 2.5-5, 5-7.5, 7.5-10, 10-15, 15+ | Patchy microbial mat |
| Clam Field | Bkgd | DR1139 | PC 56 | 36.7356; -122.0342 | 895 | 0-2.5, 2.5-5, 5-7.5, 7.5-10, 10-15, 15+ | Sand |
| Clam Field | Bkgd | DR1139 | PC 66 | 36.7365; -122.0342 | 908 | 0-2.5, 2.5-5, 5-7.5, 7.5-10, 10-15, 15+ | Sand |
| Extrovert Cliff | Seep | DR1140 | PC 64 | 36.7765; -122.0850 | 965 | 0-2.5, 2.5-5, 5-7.5, 7.5-10, 10-15 | Thick microbial mat |
| Extrovert Cliff | Seep | DR1140 | PC 44 | 36.7765; -122.0850 | 965 | 0-2.5, 2.5-5, 5-7.5, 7.5-10 | Thick microbial mat |
| Extrovert Cliff | Bkgd | DR1140 | PC 79 | 36.7765; -122.0851 | 965 | 0-2.5, 2.5-5, 5-7.5, 7.5-10, 10-15, 15+ | Sand |
| Extrovert Cliff | Bkgd | DR1140 | PC 63 | 36.7759; -122.0844 | 990 | 0-2.5, 2.5-5, 5-7.5, 7.5-10, 10-15, 15+ | Sand |

Table S2: Primers used in this study.

| Primer Name | Target Gene | Sequence (5' - 3') | Reference |
|---|---|---|---|
| mehtaFw-28 | *nifH* | GGHAARGGHGGHATHGGNAARTC | Mehta et al. (2003) |
| mehtaRv-416 | *nifH* | GGCATNGCRAANCCVCCRCANAC | Mehta et al. (2003) |

Table S3: Summary statistics for positive (mock communities) and negative controls.

| | **Mock Community** | **Negative Control** |
|---|---|---|
| **Weighted UniFrac Dissimilarity** | 0.232 ± 0.171 | - |
| **Filtered Reads** | 101881 ± 102964 | 8.27 ± 10.09 |
| ***nifH* Reads** | 33527 ± 34232 | 2.64 ± 6.20 |
| **Homolog Reads** | 68354 ± 69519 | 5.64 ± 5.57 |

Table S4: List of phyla (and classes) assigned to bona fide *nifH* genes in seep and background sediments from each of the six sites. Phyla marked with an "X" at each site if present.

| | | | **Seep** | | | | | | **Background** | | | | | |
|---|---|---|---|---|---|---|---|---|---|---|---|---|---|---|
| **Phylum as Identified by PPIT** | **Equivalent 2023 International Code of Nomenclature of Prokaryotes (ICNP) Phylum** | **Classes** | **USAM-SE** | **USAM-SW** | **USAM-NE** | **USAM-VC** | **MB-CF** | **MB-EC** | **USAM-SE** | **USAM-SW** | **USAM-NE** | **USAM-VC** | **MB-CF** | **MB-EC** |
| Euryarchaeota | Halobacteriota | Ca. Methanoliparia, Methanobacteria, Methanomicrobia, Thermoplasmata | x | x | x | x | x | x | | x | x | x | x | x |
| Acidobacteria | Acidobacteriota | Holophagae | | | x | | | | x | x | x | x | x | x |
| Actinobacteria | Actinomycetota | Actinobacteria | | | | | x | | | | | | x | |
| Bacteroidetes | Bacteroidota | Bacteroidia, Flavobacteriia | | x | x | x | x | x | x | x | x | x | x | x |
| Ca. Dadabacteria | Desulfobacterota | | | | x | x | | | | | x | x | | |
| Chlorobi | Chlorobiota | Chlorobia | x | x | x | x | x | | x | x | x | x | x | x |
| Chloroflexi | Chloroflexota | Dehalococcoidia | | | | | x | | | | | | x | |
| Cyanobacteria | Cyanobacteriota | Oscillatoriophycideae | | | x | | x | | x | | x | x | x | x |
| Deferribacteres | Deferribacterota | Deferribacteres | | | | | | | | | | x | | |
| Firmicutes | Bacillota | Bacilli, Clostridia, Negativicutes | x | x | x | x | x | x | | x | x | x | x | x |
| Fusobacteria | Fusobacteriota | Fusobacteriia | | | | | x | | x | | | | | |
| Kiritimatiellaeota | Kiritimatiellota | Kiritimatiellae | x | x | x | x | x | x | x | x | x | x | x | x |
| Lentisphaerae | Lentisphaerota | | x | x | x | x | x | x | x | x | x | x | x | x |
| Nitrospirae | Nitrospirota | Nitrospira | x | x | x | x | x | x | x | x | x | x | x | x |
| Planctomycetes | Planctomycetota | Planctomycetia | x | x | x | x | x | x | x | x | x | x | x | x |
| Proteobacteria | Pseudomonadota, Desulfobacterota* | Alpha-, Beta-, Ca. Lambda-, Delta-*, Epsilon-, Gamma-, Zeta- | x | x | x | x | x | x | x | x | x | x | x | x |
| Spirochaetes | Spirochaetota | Spirochaetia | x | x | x | x | x | | x | | x | x | x | |
| Verrucomicrobia | Verrucomicrobiota | Opitutae, Spartobacteria | x | | | x | x | | x | x | x | | x | x |

Table S5: List of phyla (and classes) assigned to bona fide *nifH* transcripts from each of the six sites. Phyla marked with an "X" at each site if present.

| Phylum as Identified by PPIT | Equivalent 2023 International Code of Nomenclature of Prokaryotes (ICNP) Phylum | Classes | USAM-SE | USAM-SW | USAM-NE | USAM-VC | MB-CF | MB-EC |
|---|---|---|---|---|---|---|---|---|
| Euryarchaeota | Halobacteriota | Ca. Methanoliparia, Methanomicrobia | x | x | x | x | | |
| Firmicutes | Bacillota | Clostridia | x | x | x | x | | |
| Proteobacteria | Desulfobacterota | Deltaproteobacteria | x | x | x | x | | x |

### Supplementary Figures

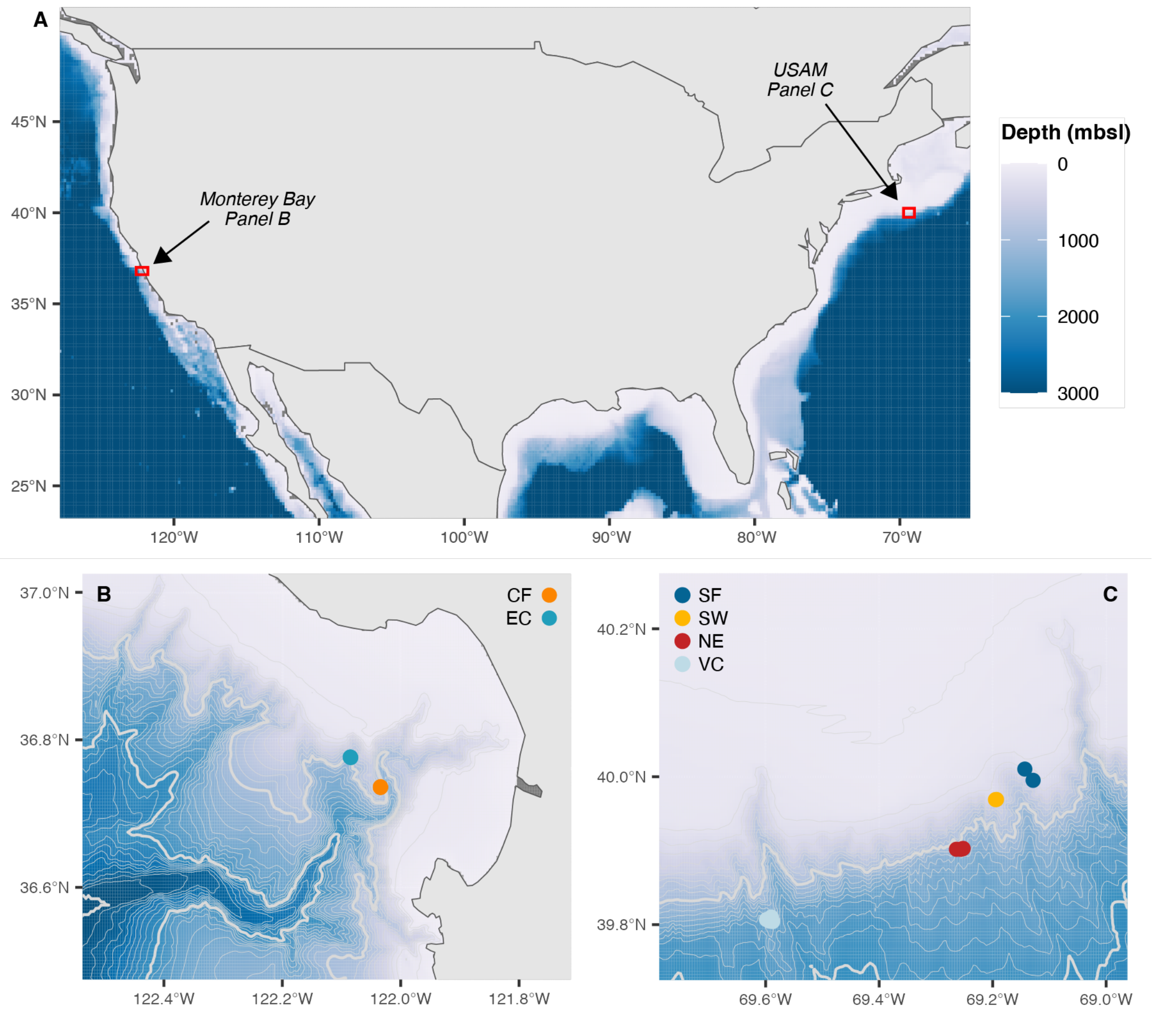


Fig S1: Location of both study regions – Monterey Bay (MB) and the northern U.S. Atlantic Margin (USAM) – outlined in red boxes (A). Cold seep sediment sampling sites in Monterey Bay (enlarged in panel B) are Clam Field (CF – 895-909 mbsl) and Extrovert Cliff (EC – 965-990 mbsl); sampling sites along the USAM (enlarged in panel C) are Shallop Canyon East (SE – 335-366 mbsl), Shallop Canyon West (SW – 390-395 mbsl), New England (NE – 1,130-1,252 mbsl), and Veatch Canyon (VC – 1,407-1,545 mbsl). Thin grey contour lines every 100 mbsl, thick grey contour lines every 1000 mbsl. Any depths over 3,000 mbsl are represented by the same dark blue color. Data are adapted from Semler et al. (2022; USAM) and Semler & Dekas (2024, preprint; MB).

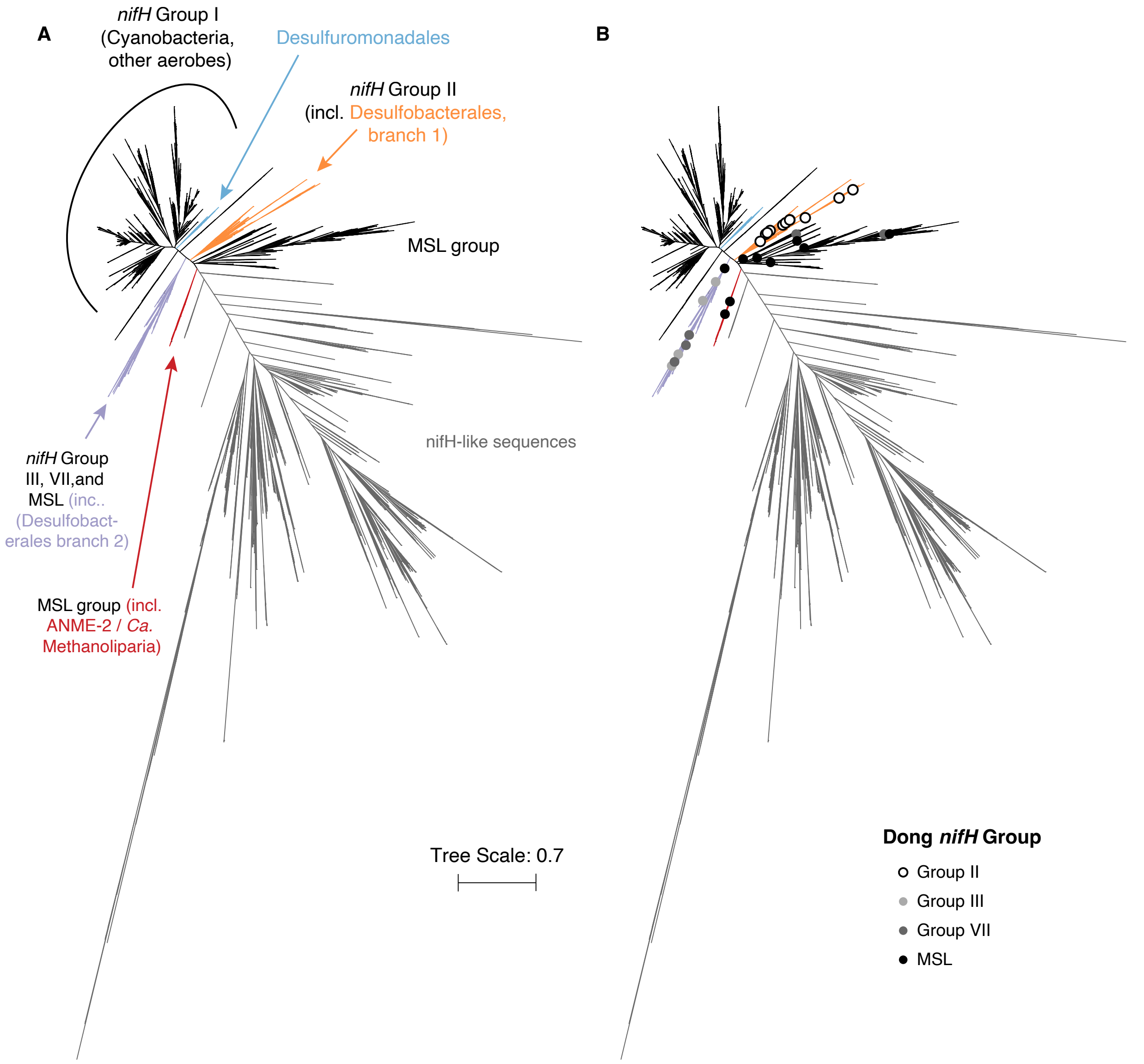


Fig. S2: Annotated *nifH* reference tree (A) used as an input for the taxonomic assignment program PPIT (Kapili and Dekas, 2020), with *nifH* sequences in MAGs from Dong et al. (2022) overlaid and labeled according to their *nifH* group designations (B). Scale bar shows the expected number of nucleotide substitutions per site.

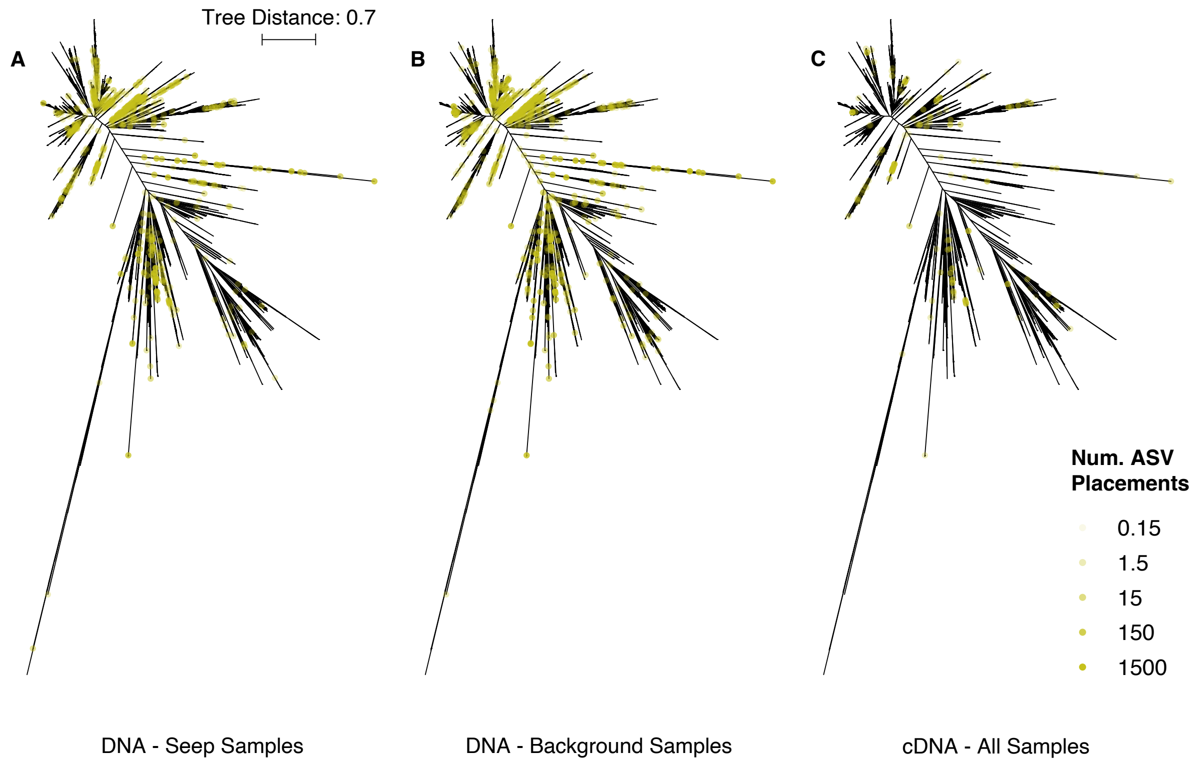


Fig. S3: Annotated *nifH* reference tree used as an input for the taxonomic assignment program PPIT (Kapili and Dekas, 2020). Location of major *nifH* groups labelled, and branches with taxa relevant to our dataset are colored. *nifH* gene ASV placements from seep samples (A) and background samples (B), and *nifH* transcript ASV placements from all samples (C) are marked with black circles; darker colors indicate more placements. Scale bar shows the expected number of nucleotide substitutions per site.

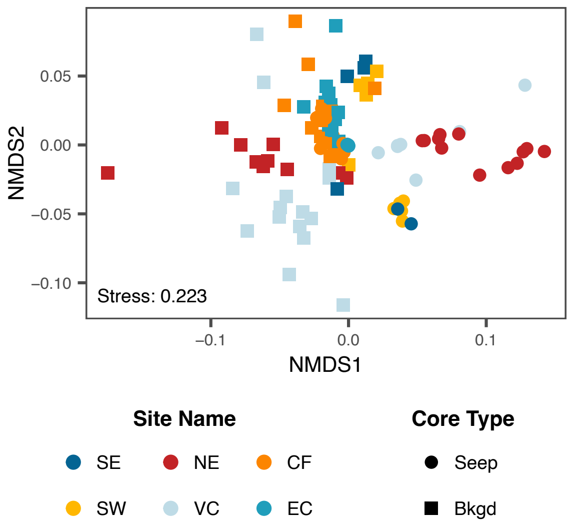


Fig S4: Non-metric multi-dimensional scaling (NMDS) plot of seep and background samples. Analysis was based on a weighted UniFrac distance metric and was inferred by *nifH* gene sequencing. *nifH*-like sequences were removed prior to analysis.

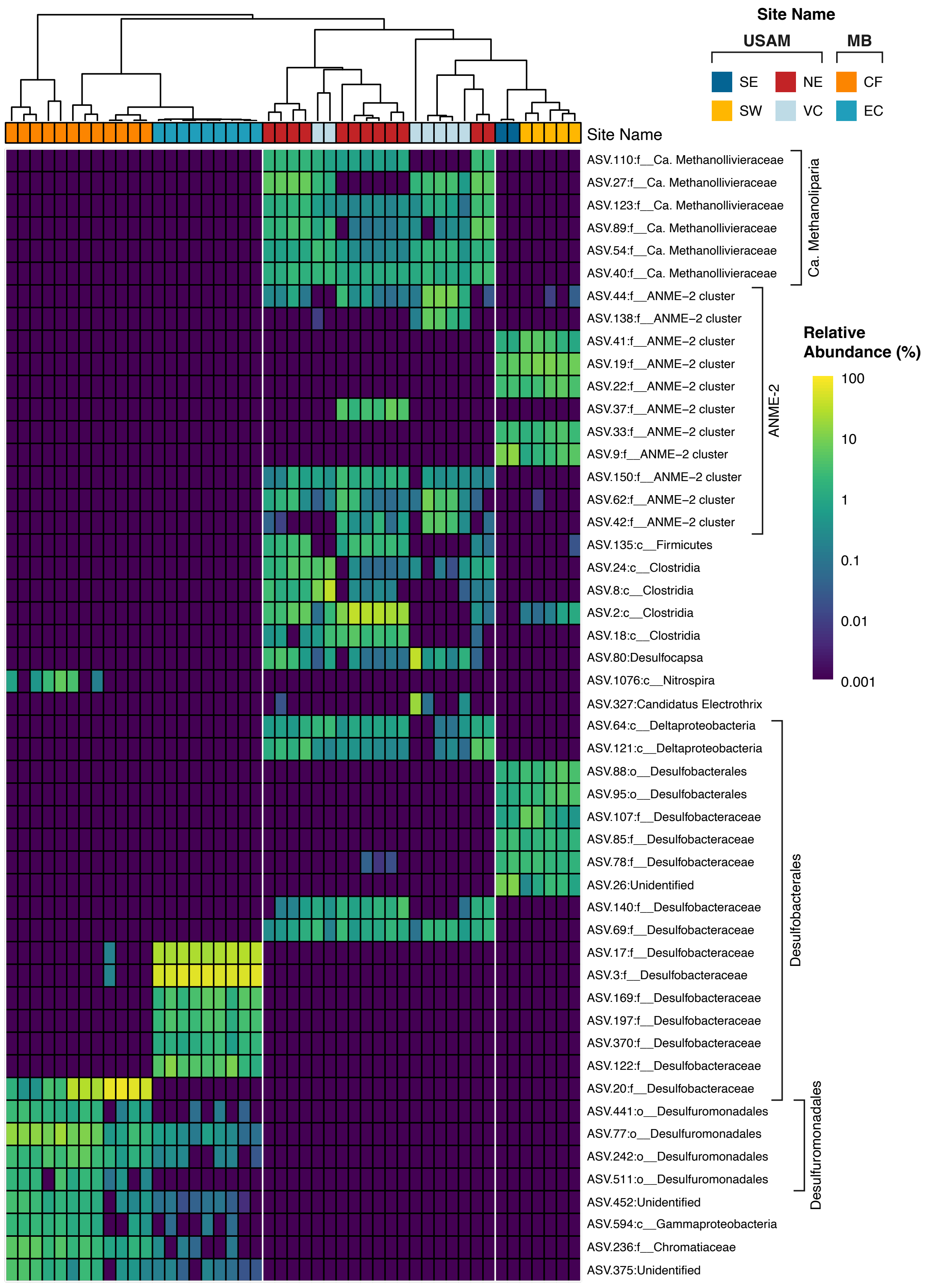
Site Name
USAM
MB
SE
NE
CF
SW
VC
EC
Site Name
Relative Abundance (%)
100
10
1
0.1
0.01
0.001
ASV.110:f__Ca. Methanollivieraceae
ASV.27:f__Ca. Methanollivieraceae
ASV.123:f__Ca. Methanollivieraceae
ASV.89:f__Ca. Methanollivieraceae
ASV.54:f__Ca. Methanollivieraceae
ASV.40:f__Ca. Methanollivieraceae
ASV.44:f__ANME−2 cluster
ASV.138:f__ANME−2 cluster
ASV.41:f__ANME−2 cluster
ASV.19:f__ANME−2 cluster
ASV.22:f__ANME−2 cluster
ASV.37:f__ANME−2 cluster
ASV.33:f__ANME−2 cluster
ASV.9:f__ANME−2 cluster
ASV.150:f__ANME−2 cluster
ASV.62:f__ANME−2 cluster
ASV.42:f__ANME−2 cluster
ASV.135:c__Firmicutes
ASV.24:c__Clostridia
ASV.8:c__Clostridia
ASV.2:c__Clostridia
ASV.18:c__Clostridia
ASV.80:Desulfocapsa
ASV.1076:c__Nitrospira
ASV.327:Candidatus Electrothrix
ASV.64:c__Deltaproteobacteria
ASV.121:c__Deltaproteobacteria
ASV.88:o__Desulfobacterales
ASV.95:o__Desulfobacterales
ASV.107:f__Desulfobacteraceae
ASV.85:f__Desulfobacteraceae
ASV.78:f__Desulfobacteraceae
ASV.26:Unidentified
ASV.140:f__Desulfobacteraceae
ASV.69:f__Desulfobacteraceae
ASV.17:f__Desulfobacteraceae
ASV.3:f__Desulfobacteraceae
ASV.169:f__Desulfobacteraceae
ASV.197:f__Desulfobacteraceae
ASV.370:f__Desulfobacteraceae
ASV.122:f__Desulfobacteraceae
ASV.20:f__Desulfobacteraceae
ASV.441:o__Desulfuromonadales
ASV.77:o__Desulfuromonadales
ASV.242:o__Desulfuromonadales
ASV.511:o__Desulfuromonadales
ASV.452:Unidentified
ASV.594:c__Gammaproteobacteria
ASV.236:f__Chromatiaceae
ASV.375:Unidentified
Ca. Methanoliparia
ANME-2
Desulfobacterales
Desulfuromonadales

Fig. S5 (previous page): Relative abundance (%) of the 50 most abundant *nifH* ASVs in all samples after removing *nifH*-like sequences. Samples clustered by the weighted UniFrac distance between samples.

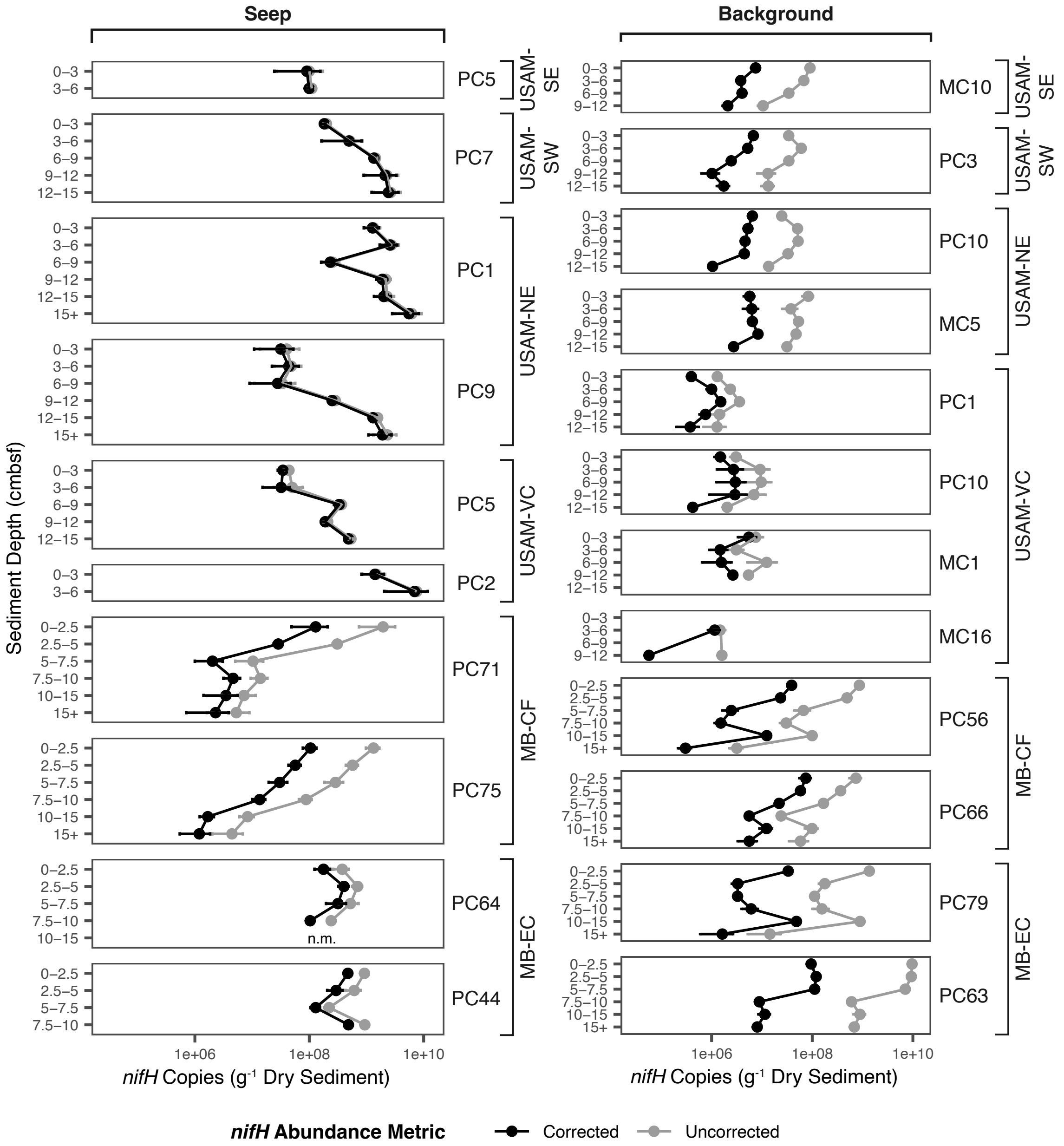


Fig. S6: Uncorrected (in grey) and corrected (in black) *nifH* gene copies (gram dry sediment$^{-1}$) with sediment depth in all seep and background cores. Error bars represent the standard deviation of triplicate measurements. Missing values indicate that *nifH* was below the limit of detection for qPCR, with the exception of the MB-EC PC64 sample from 12-15 cmbsf, which was not measured due to sample evaporation.

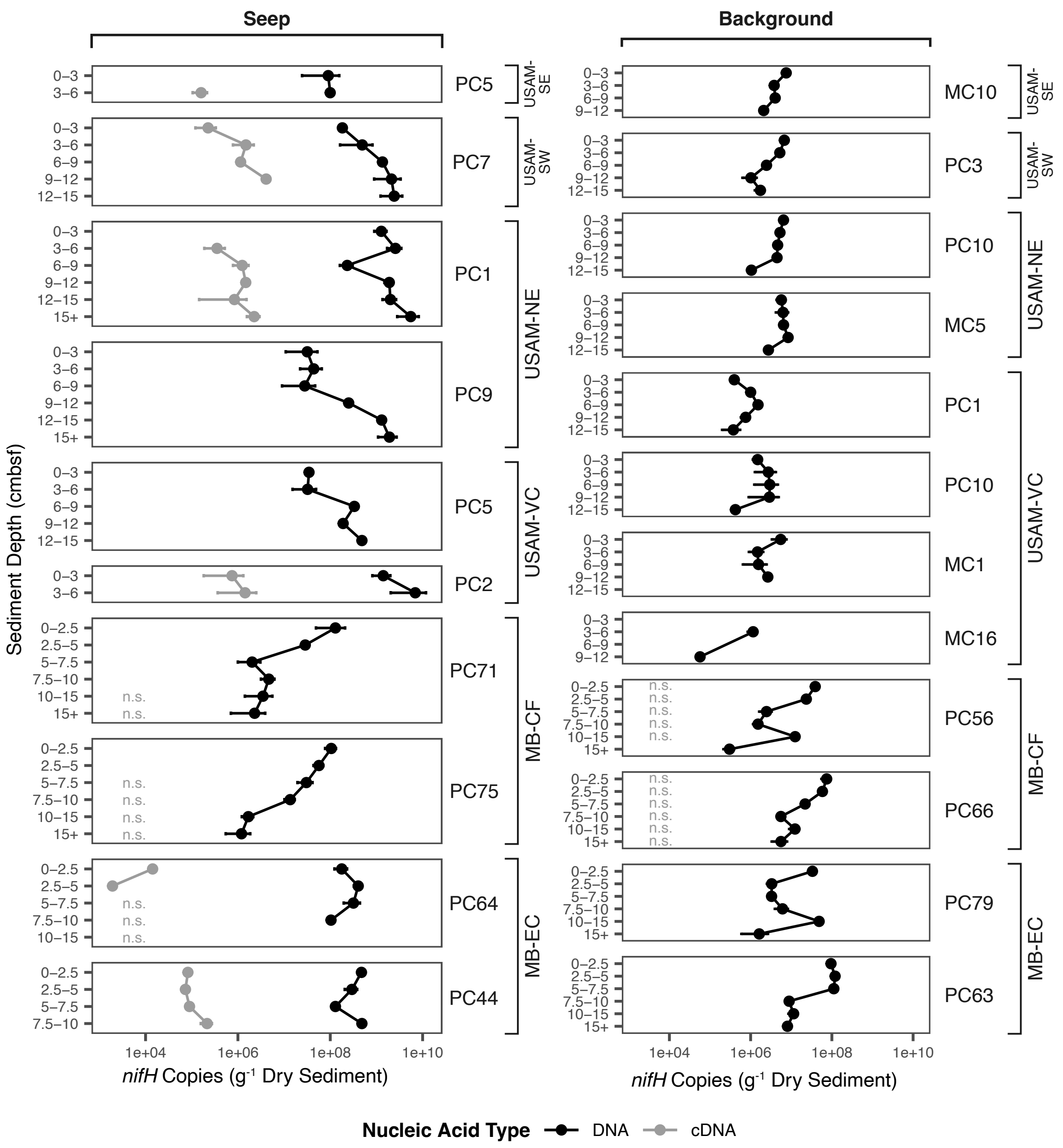


Fig. S7: Abundance of *nifH* genes (DNA) and transcripts (cDNA) with sediment depth in all seep and background cores. Values were corrected by adjusting raw abundances downward by the percentage of sequences determined to be *nifH*-like by coincident sequencing data. Error bars represent the standard deviation of triplicate measurements. Missing values indicate that *nifH* was below the limit of detection for qPCR, with the exception of the Extrovert Cliff PC64 DNA sample from 12-15 cmbsf, which was not measured due to sample evaporation.

n.s. – sample was not sequenced (no observable *nifH* amplification) and, despite a successful qPCR measurement, has no corresponding sequencing data available to correct raw qPCR measurement

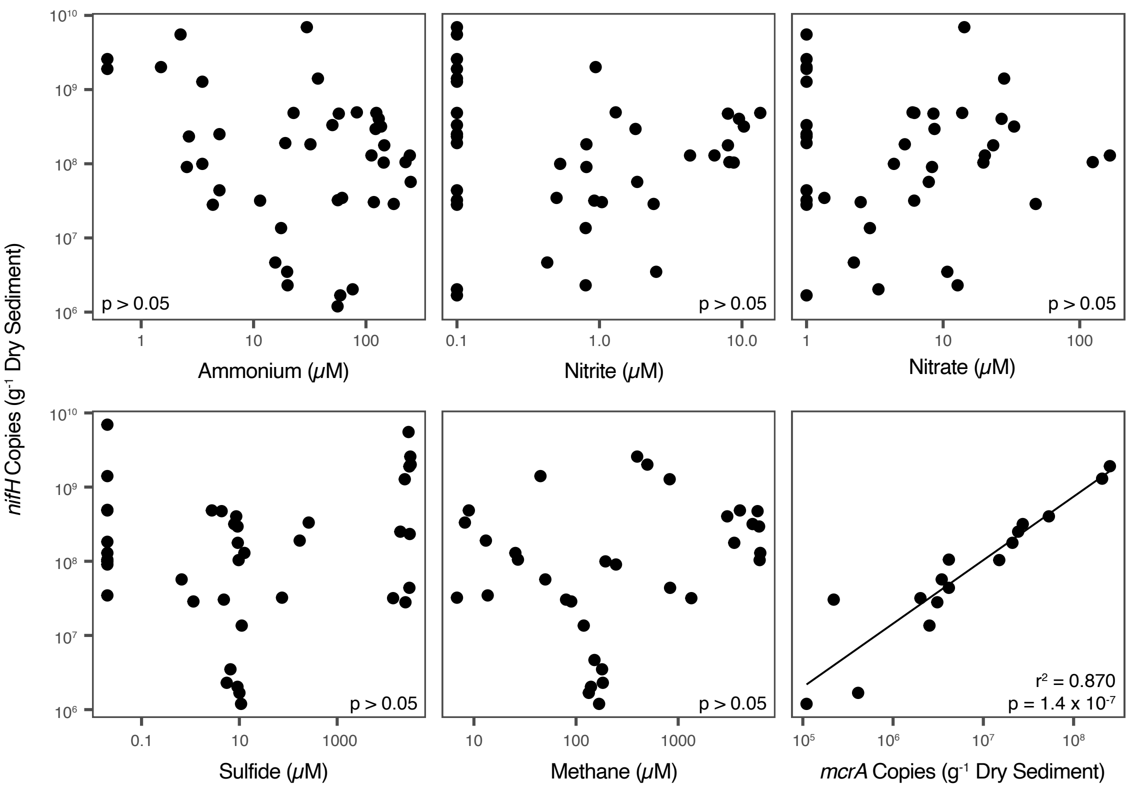


Fig. S8: Corrected *nifH* gene abundances in seep samples compared to concentrations of ammonium, nitrite, nitrate, sulfide, and methane, and to *mcrA* gene abundances. Significant relationships (p-value < 0.05) shown with solid trendlines; those with no significance (p-value > 0.05) shown with no trendlines.
